# Iridescence in pterosaur pycnofibers and the evolution of integumentary coloration

**DOI:** 10.64898/2026.05.06.723168

**Authors:** Zelin Wu, Liliana D’ Alba, Chang-Fu Zhou, Julia A. Clarke, Jinhua Li, Matthew D. Shawkey, Quanguo Li

## Abstract

The bodies of pterosaurs, the first flying vertebrates, are covered with integumentary filaments (pycnofibres) thought to be homologous to feathers in dinosaurs, but their coloration remains unknown. Here, we report a layered internal arrangement of melanosomes containing a photonic nanostructure within the monofilaments in a previously undescribed specimen of tapejarid pterosaur *Sinopterus dongi* from the Early Cretaceous Jehol Biota. Optical simulations showed that this structure reflects green to magenta iridescent coloration, confirming the presence of melanosome-based iridescent coloration previously thought to be unique to birds. This finding deepens our understanding of structure/color gamut relationships in amniotes, while supporting further shared characteristics associated with derived genetic and regulatory shifts in archosaurs.

## Main Text

The physical appearance of extinct animals has long been one of the most contentious issues in paleontology. Previously thought to be impossible, the assignment of melanin-based colors to fossilized integument was enabled with the discovery of melanosomes in exceptionally preserved fossils in 2008 (*1–4*). Melanosomes are organelles that produce and store melanins, and are involved in many functions including color formation (*5, 6*). Nanoscale organization of melanosomes can generate iridescent structural color through coherent scattering of light (*7*), responsible for some of the most brilliant colors of birds. Melanosomes (or their imprints) have been widely reported in the soft tissue of vertebrate fossils including dinosaurs and pterosaurs (*8*). Evidence for structural color has also been detected in the feathers of paravian theropods *Microraptor* (*1*), *Caihong* (*9*), and enantiornithine bird fossils (*10*), but is not known in the integumentary structures of vertebrates other than Paraves (a clade of maniraptoran theropods including Avialae and close relatives).

“Hair-like” integumentary structures (termed pycnofibres) (*11*) are found in well-preserved pterosaur fossils. Although usually present as unbranched monofilaments, different types of branched pycnofibres from pterosaurs have been recently revealed (*12, 13*). Pycnofibres are proposed to be homologous with avian feathers in terms of branching patterns, melanosome diversity and genetic underpinnings (*12–15*), but their internal structure and similarities to avian feathers remain unknown. These uncertainties hinder us from extending the melanosome morphology-coloration correlations from paravian dinosaurs to pterosaurs (*1*). Therefore, exploring the internal structure of pycnofibres is critical to reconstruct pterosaur coloration and the evolution of integumentary structure in archosaurians.

In this study we report a previously undescribed specimen of tapejarid pterosaur - *Sinopterus dongi* from the Lower Cretaceous Jiufotang Formation (Lamadong locality, Jianchang county, Liaoning Province, China; Fig. 1). It is an incomplete skeleton with extensive soft tissues, including preserved remains of pycnofibres, skin, claw sheaths and wing membrane (Fig. 1, fig. S1, S2I, J, see supplementary text for detailed description). We used micro X-ray fluorescence spectrometry (Micro-XRF), scanning electron microscopy (SEM), dual-beam focused ion-beam scanning electron microscopy (FIB-SEM), energy-dispersive X-ray spectrometer (EDS) and time of flight secondary ion mass spectrometry (ToF-SIMS) to explore the chemical composition and microstructure of pycnofibres and other soft tissues. We found a photonic structure as well as complex internal organization of melanosomes in monofilaments of *Sinopterus dongi*, and reconstructed its iridescent color by finite-difference time-domain (FDTD) analyses. The diversity of melanosomes, and their organization in monofilaments suggests a complex pattern of evolution of structural color present today in avian barbs and barbules.

**Fig. 1.**
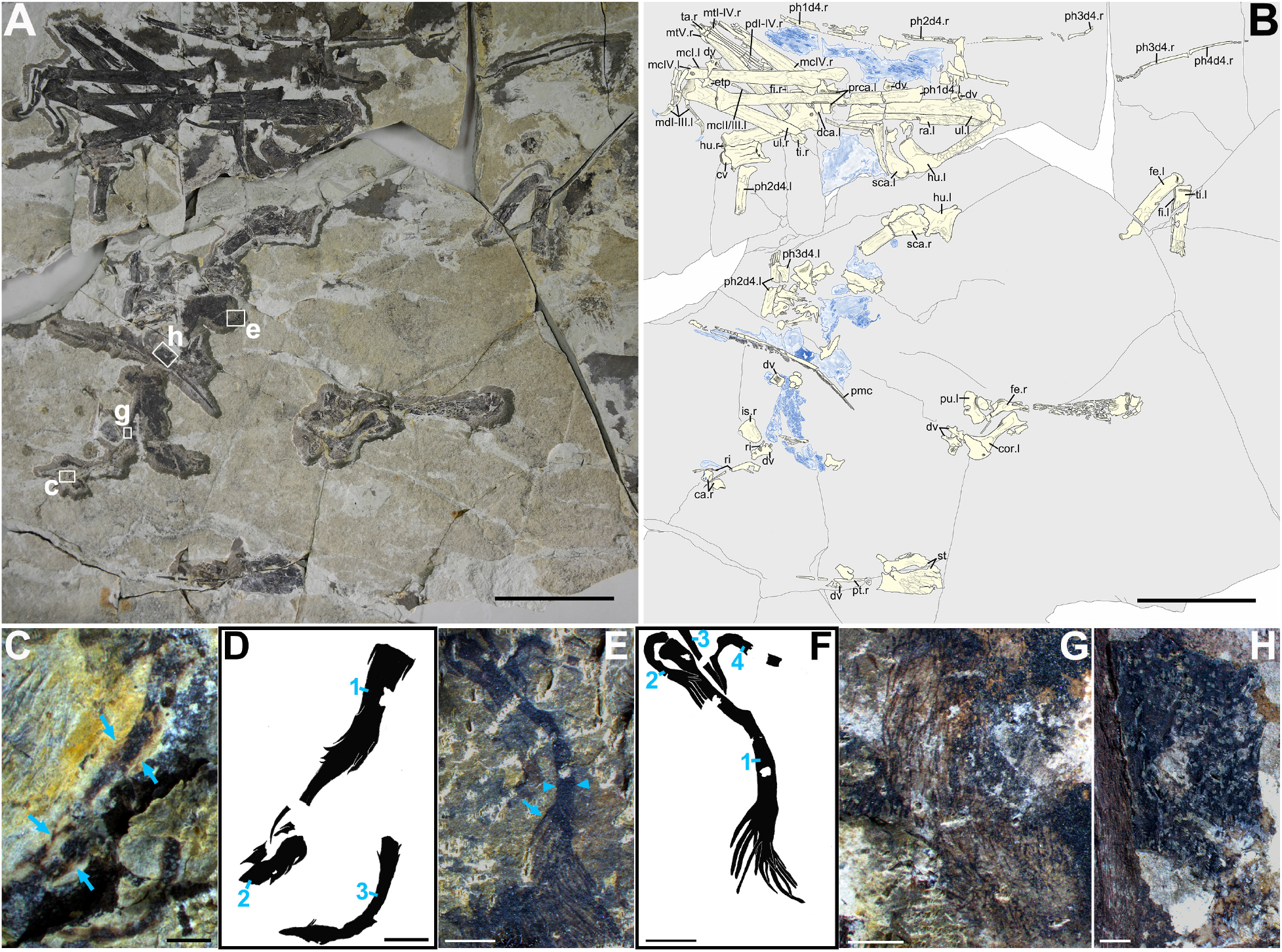
Photograph, line drawing of *Sinopterus dongi* CUGB-P2201 and details of soft tissues. (**A**), Photograph of CUGB-P2201. (**B**), Line drawing of CUGB-P2201, skeleton and soft tissues are marked in yellow and blue respectively. (**C** to **H**), Details of soft tissues. (**C**),”Feather-like” pycnofibres. Arrows: branches on either side. (**D**), Outline silhouette of “feather-like” pycnofibres (marked by 1-3). (**E**),”Brush-like” pycnofibres. Arrow, distal branches; Arrowheads, the thick and elongated proximal shaft. (**F**), Outline silhouette of “brush-like” pycnofibres (marked by 1-4). Numbers in (D) and (F) represent different individuals of the same type. (**G**), Monofilaments. (**H**), Fossilized skin remnant located on the ventral side of the premaxillary crest. Abbreviations: ca, carpal; cor, coracoid; cv, cervical vertebra; dca, distal carpal; dv, dorsal vertebra; etp, extensor tendon process; fe, femur; fi, fibula; hu, humerus; is, ischium; mc I-IV, metacarpal I-IV; md I-III, manual digits I-III; mt I-V, metatarsal I-V; pd I-V, pedal digits I-V; ph1d4-ph4d4, first-fourth phalanx of manual digit IV; pmc, premaxillary crest; prca, proximal carpal; pt, pteroid; pu, pubis; ra, radius; ri, rib; sca, scapula; st, sternum; ta, tarsal; ti, tibia; ul, ulna; l, left; r, right. Scale bars, 10cm (A, B); 2 mm (E-H); 1 mm (C-D).

### Elemental composition of soft tissues

Micro-XRF shows clear differences in the chemical elements of the rocky matrix, skeleton and soft tissues (fig. S3). The elements, K, Si, Fe, Al and Mn, are distributed in the rocky matrix; P, Ca, S, Sr and Y are associated with the skeleton; S, Cu, Ni and Ti are rich in the pycnofibres and fossilized skin remnants on the ventral margin of the premaxillary crest, of which Cu enrichment is highly correlated with soft tissues (fig. S4C-F). Cu, Ti, Ni and S show ribbon-like patterns on the crest skin remnant (fig. S4C-F, insets), and are enriched in three striped regions, most obviously immediately adjacent to the crest. The specific distribution of elements indicates different composition among soft tissues. The FIB results showed considerable enrichment of microbodies in the element-enriched stripe regions, whereas the element-depleted regions had very few or no microbodies (fig. S4G-I). These microbodies should represent melanosomes based on the uniform internal electron density and the ion peaks associated with eumelanin revealed by ToF-SIMS (fig. S4H, see supplementary text for more details). Regional differences of elemental distribution and melanosomes concentrations suggest the skin near premaxillary crest of *Sinopterus dongi* (and even the adjacent soft tissue crest) had a melanin-based stripe pattern, similar to that proposed for *Pterorhynchus wellnhoferi* (*16*).

### Photonic structure and internal organization of melanosomes in pycnofibres

Five types of pycnofibre have been previously reported in pterosaurs (*12, 13*), and three of these can be identified here: monofilaments (no branching) (Fig. 1G), “brush-like” (terminal branching) (Fig. 1E) and “feather-like” (branching along both sides of the axis) pycnofibres (Fig. 1C). The monofilaments of this pterosaur are most abundant and are therefore crucial to their appearance. We used FIB to cut 16 monofilaments with no skin remnant influence (fig. S5A-Eb) into 43 sections, and found that their interiors contained numerous nanoscale microbodies encapsulated in a matrix (Fig. 2C,F,I, fig. S6-S8). The morphology, arrangement and chemical composition of these microbodies support that they are melanosomes, and the matrix should represent a mineralized product of original composition (fig. S9, see supplementary text for more details).

**Fig. 2.**
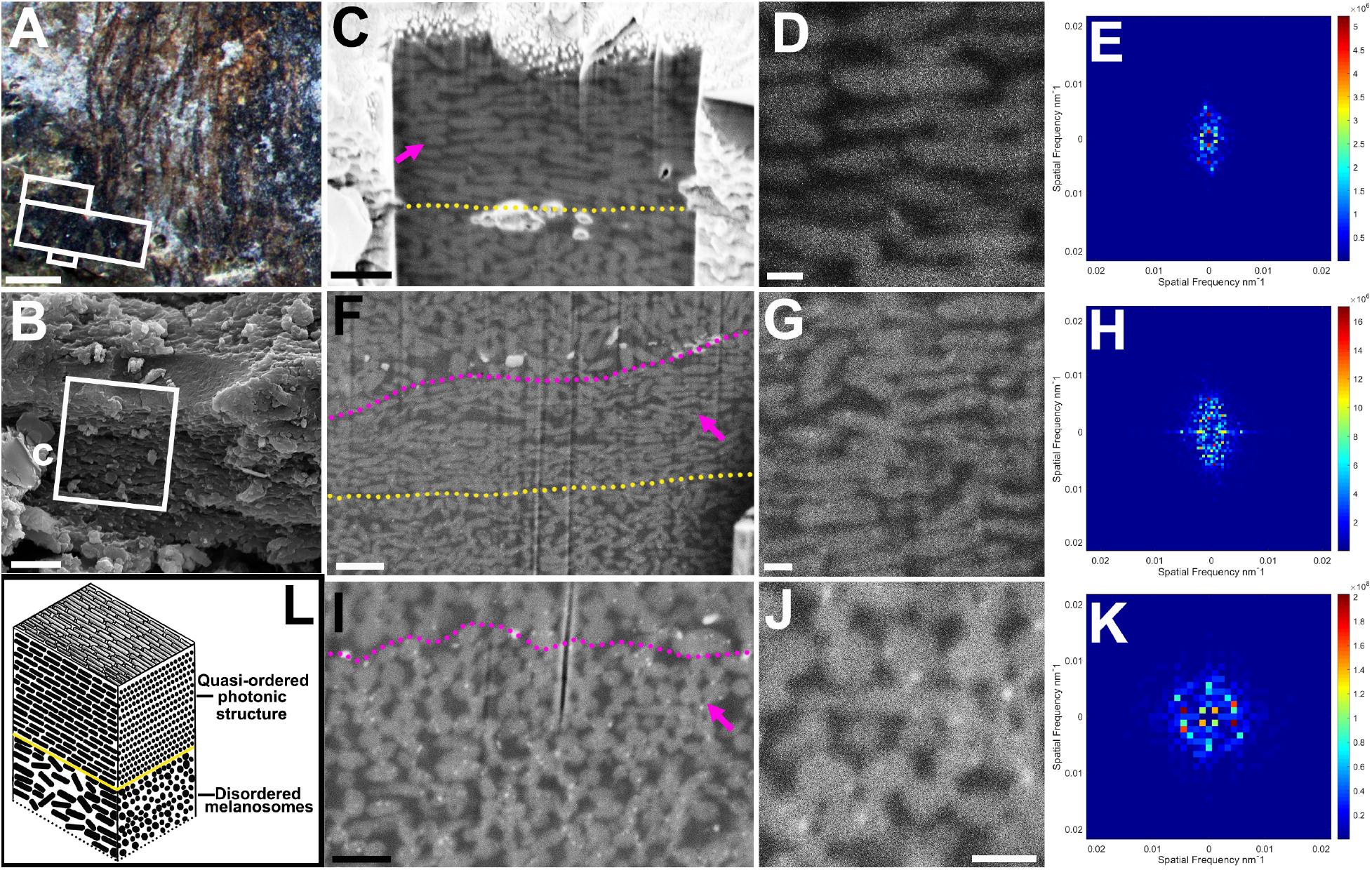
Photonic structure of melanosomes in monofilaments. (**A**). Sample location (white boxes). (**B**), Monofilament at position M1ls1, showing a smooth surface. (**C** to **H**), Longitudinal sections and 2-D Fourier power spectra of the monofilament at position M1ls1 (**C** to **E**) and M2csls4 (**F** to **H**), showing highly ordered arrangement of melanosomes in the vertical direction. (**I** to **K**), Cross section (**I** to **J**) of the monofilament at position M2csls4 corresponding to (F) and its 2-D Fourier power spectra (**K**), showing a nearly hexagonal photonic structure. Violet arrows in (C, F, I) point to the region used for 2-D Fourier analysis, (D, G, J) are close-up of these regions. The yellow dashed lines in (C, F) mark the boundary between the photonic structure and disordered melanosomes below it. Note the monofilament corresponding to (F, I) has undergone staggered slip, resulting in the melanosomes at the bottom superimposed on the top (violet dashed lines). (**L**), Reconstruction model of photonic structure, melanosomes are not drawn to scale. Secondary electron images (B to D); Backscattered electron images (F, G, I, J). Scale bars, 1mm (A); 2 μm (B); 1 μm (C, F); 500 nm (I); 200 nm (D, G, J).

We found direct evidence for iridescent structural color in 2 of 16 monofilaments. The section at position M1ls1 shows 15 layers of rod-like melanosomes at the top. The average diameter of melanosomes is 149.80 ± 12.08 nm (n=14), organized head to tail and parallel to one another, in highly ordered arrays (violet arrow in Fig. 2C). These structures should be located on the outermost side of monofilaments because position M1ls1 has a naturally smooth surface (Fig. 2B), similar to extant iridescent feathers (*7*). The same structure was also found at position M2csls4 (violet arrow in Fig. 2F). The section here shows 16 layers of regularly arranged rod-like melanosomes. Unlike M1ls1, the melanosome array here was overlaid by some melanosomes with larger size at an angle of 10° (violet dashed line in Fig. 2F, I), these overlays represent taphonomic artifacts rather than the original structure (supplementary text). The ordered melanosomes from both M1ls1 and M2csls4 have similar thin diameters (147.90 ± 13.77 nm, n=55) as well as high aspect ratios (4.11 ± 0.34, n=55, Table 1). The cross sections corresponding to position M2csls4 show that some melanosomes still retained the original arrangement: they are aligned in a nearly hexagonal array perpendicular to the cross section (Fig. 2I, fig. S10A-F), resembling the iridescent wing patch of extant ducks (*17*). The total thickness of the ordered melanosome layer is nearly 2. 5 μm.

**Table 1.**
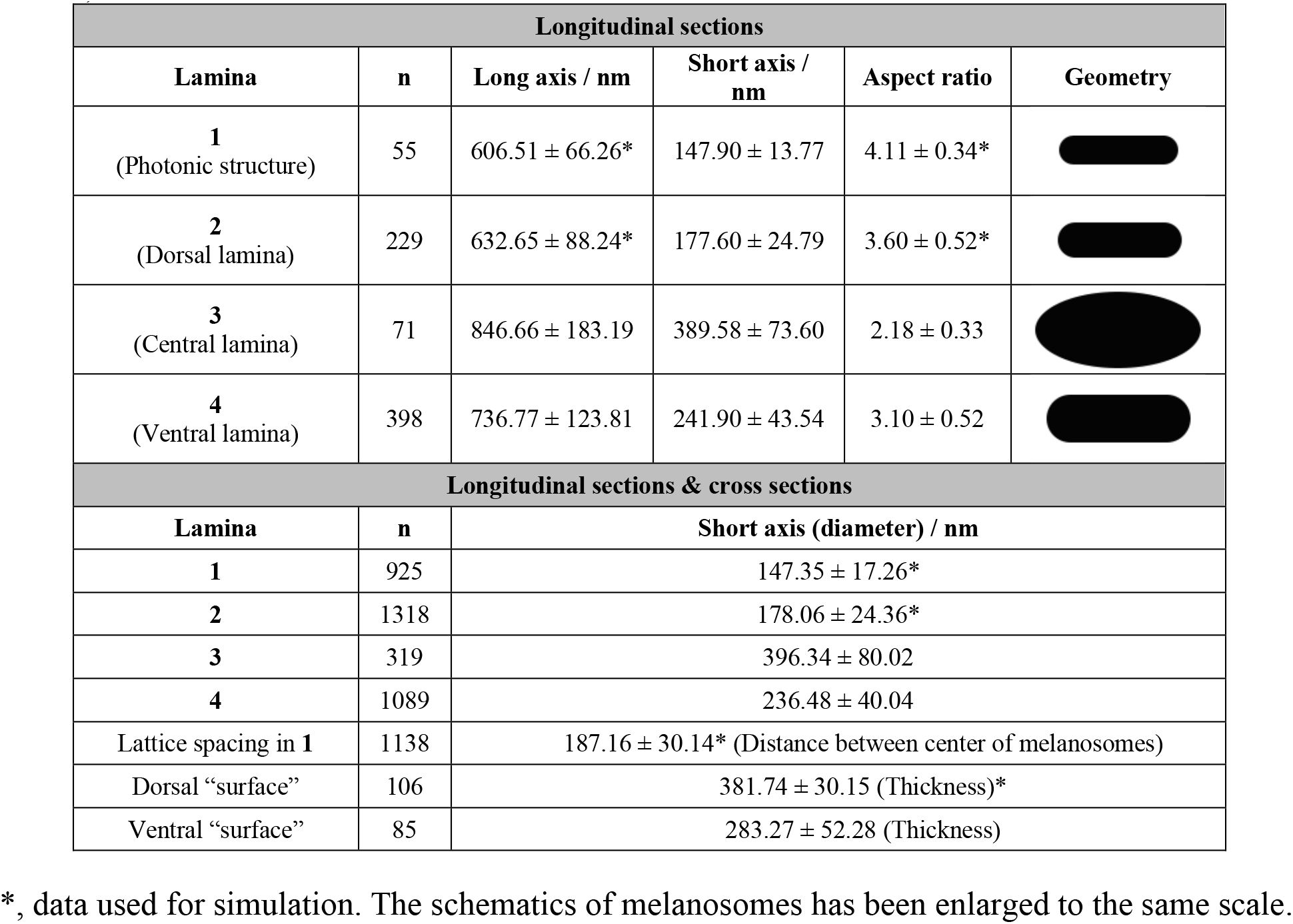
Morphological data of melanosomes (mean ± standard deviation) measured from longitudinal sections and cross sections of monofilaments in *Sinopterus dongi* (CUGB-P2201).

| Longitudinal sections |  |  |  |  |  |
| --- | --- | --- | --- | --- | --- |
| Lamina | n | Long axis / nm | Short axis / nm | Aspect ratio | Geometry |
| 1<br>(Photonic structure) | 55 | 606.51 ± 66.26* | 147.90 ± 13.77 | 4.11 ± 0.34* | |
| 2<br>(Dorsal lamina) | 229 | 632.65 ± 88.24* | 177.60 ± 24.79 | 3.60 ± 0.52* | |
| 3<br>(Central lamina) | 71 | 846.66 ± 183.19 | 389.58 ± 73.60 | 2.18 ± 0.33 | |
| 4<br>(Ventral lamina) | 398 | 736.77 ± 123.81 | 241.90 ± 43.54 | 3.10 ± 0.52 | |
| Longitudinal sections & cross sections |  |  |  |  |  |
| Lamina | n | Short axis (diameter) / nm |  |  |  |
| 1 | 925 | 147.35 ± 17.26* | | | |
| 2 | 1318 | 178.06 ± 24.36* | | | |
| 3 | 319 | 396.34 ± 80.02 | | | |
| 4 | 1089 | 236.48 ± 40.04 | | | |
| Lattice spacing in 1 | 1138 | 187.16 ± 30.14* (Distance between center of melanosomes) | | | |
| Dorsal “surface” | 106 | 381.74 ± 30.15 (Thickness)* | | | |
| Ventral “surface” | 85 | 283.27 ± 52.28 (Thickness) | | | |

\*, data used for simulation. The schematics of melanosomes has been enlarged to the same scale.

To further investigate the regularity of the melanosome arrangement, we performed 2-D Fourier analysis on undistorted regions. The results of the longitudinal sections at position M1ls1 and M2csls4 both showed symmetric regions of high values above and below the origin (Fig. 2E,H), which suggests the structure is ordered and periodically varies in the vertical direction. In cross section, the 2-D Fourier analysis of the region at position M2csls4 showed a symmetrical hexagonal pattern (Fig. 2K, fig. S10G-I), suggesting the presence of a two-dimensional hexagonal photonic structure within monofilaments, most similar to that observed in iridescent wing patch of extant ducks (*17*). Here we define the ordered photonic structure as “Lamina-1” (Table 1). To our knowledge, this represents the first report of pterosaur melanosomes with structuring similar to “thin melanin layers” in avian iridescent feathers (i.e., melanin layer within melanosomes <190 nm, fig. S11D) (*27*), validating our inference.

In both section M1ls1 and M2csls4, there are disordered rod-shaped melanosomes distributed below the photonic structure (the melanosomes below the yellow dashed lines in Fig. 2C,F, 175.06 ± 19.62 nm, n=35), which suggests the presence of other melanosomes in monofilaments other than the photonic structure. We reconstructed the internal structure of a monofilament based on the hierarchical organization of melanosomes found in different sections. A complete monofilament contains the photonic structure and three additional laminae of melanosomes with different sizes and morphologies (fig. S6-S8,S11-S12, see supplementary text for a detailed description of the internal structure in monofilaments).

### Optical simulation of pycnofibre color

To estimate the reflective color properties of the nanostructures in the fossil, we performed finite-difference time-domain (FDTD) analyses. Since the exact nature of the matrix in pterosaur monofilaments is unknown, we modeled the fossil nanostructures with the keratin matrix that was found in extant birds. Simulations estimated that ordered melanosome nanostructures produced reflectance curves with peaks in the visible wavelengths that change with angle of incident light. A three-dimensional model of idealized hexagonal melanosome photonic crystal constructed based on fossil data produced a greenish color at 0° incident angle with a primary peak that slightly shifted towards shorter wavelengths as the incident angle increases, accompanied by a decrease in reflectance, and a blue hue at 30°. Then the primary peak shifted to an intense dark red wavelength (664 nm) at 40° (Fig. 3C). When a potential cortex surface is considered (see supplementary text for a discussion of potential surface in monofilaments), the reflection peaks exhibited minor shifts at 0°-30°, and a more intense dark red peak at 40° and 50° (Fig. 3A), indicating that the keratin cortex primarily affect brightness and contrast. When both a potential cortex surface and previously proposed taphonomic shrinkage of melanosomes (*18, 19*) (10%, see supplementary text for an estimate of the extent of melanosome shrinkage) are modeled, the reflection peak remain green at 0°-10° and intense dark red at 40°, but a less saturated red color at 20°, 30° and 50° (fig. S13A). To assess the effect of model ordering on the spectra, we also simulated two-dimensional models directly based on cross sections of monofilament, showing that even the nanostructure as preserved produces yellow-greenish to dark magenta iridescent colors regardless of the presence (Fig. 3B) or absence of a cortex (Fig. 3D, fig. S13B-D). These results suggest that structural color production is robust to presence or absence of a keratinous cortex surface, taphonomic shrinkage of melanosomes, or model ordering, and that the monofilaments of *Sinopterus dongi* produce iridescent colors , with most likely green/blue to dark magenta iridescent hues (Fig. 4). While our modeling has limitations, including assumed refractive index values, it strongly suggests that, at minimum, these pycnofibres contain an optically active nanostructure.

**Fig. 3.**
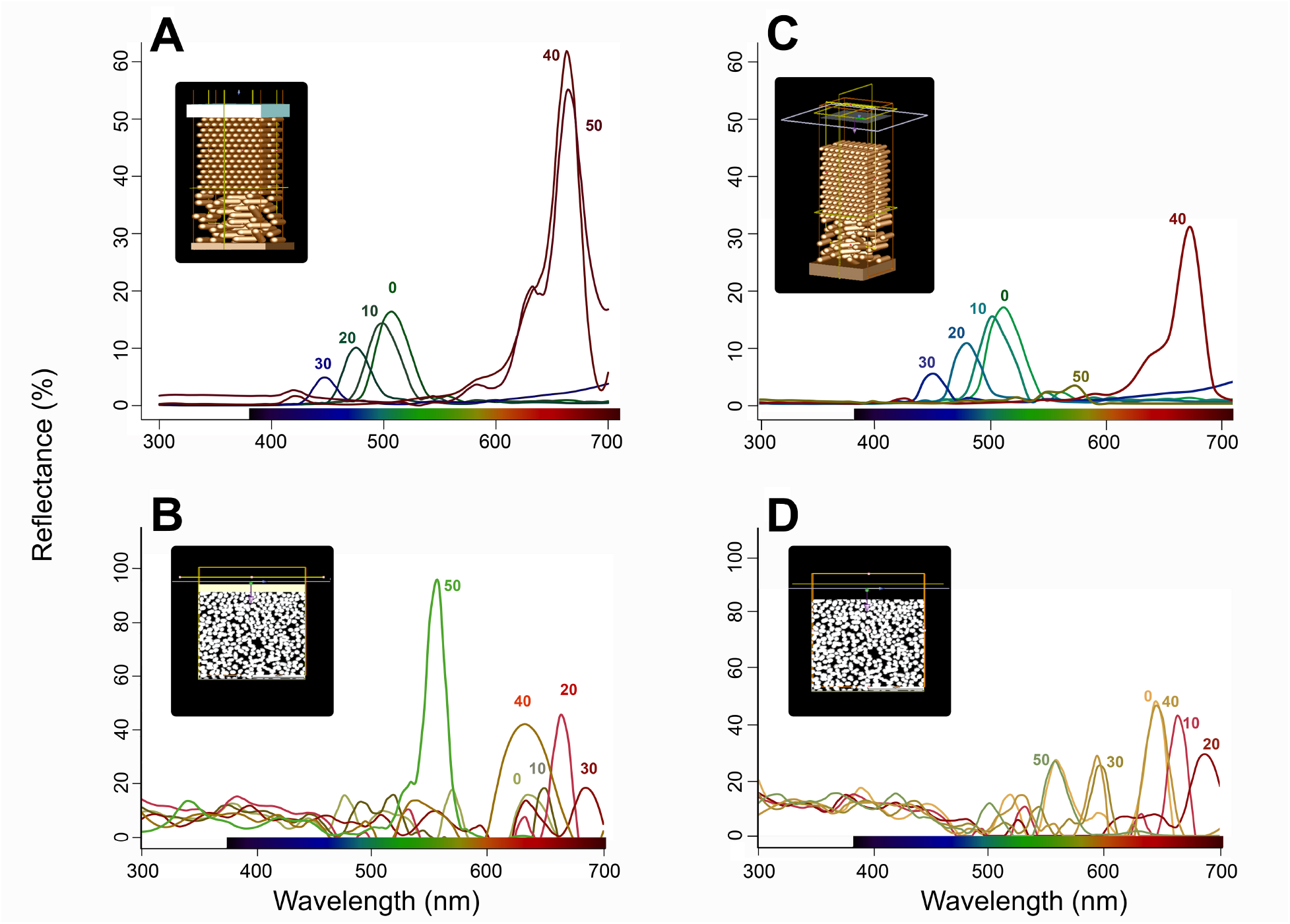
Simulated angle-dependent reflectance from models of idealized and realistic melanosome structures in monofilaments. (**A**), Idealized model with putative keratin cortex, normal size melanosomes and keratin background. (**B**), Realistic model based on the fossil section shown in Fig. 2I, modeled with keratin background and thin keratin cortex. (**C**), Idealized model without keratin cortex, with normal size melanosomes and keratin background. (**D**), Realistic models based on the fossil section shown in Fig. 2I, modeled with keratin background but no keratin cortex.

**Fig. 4.**
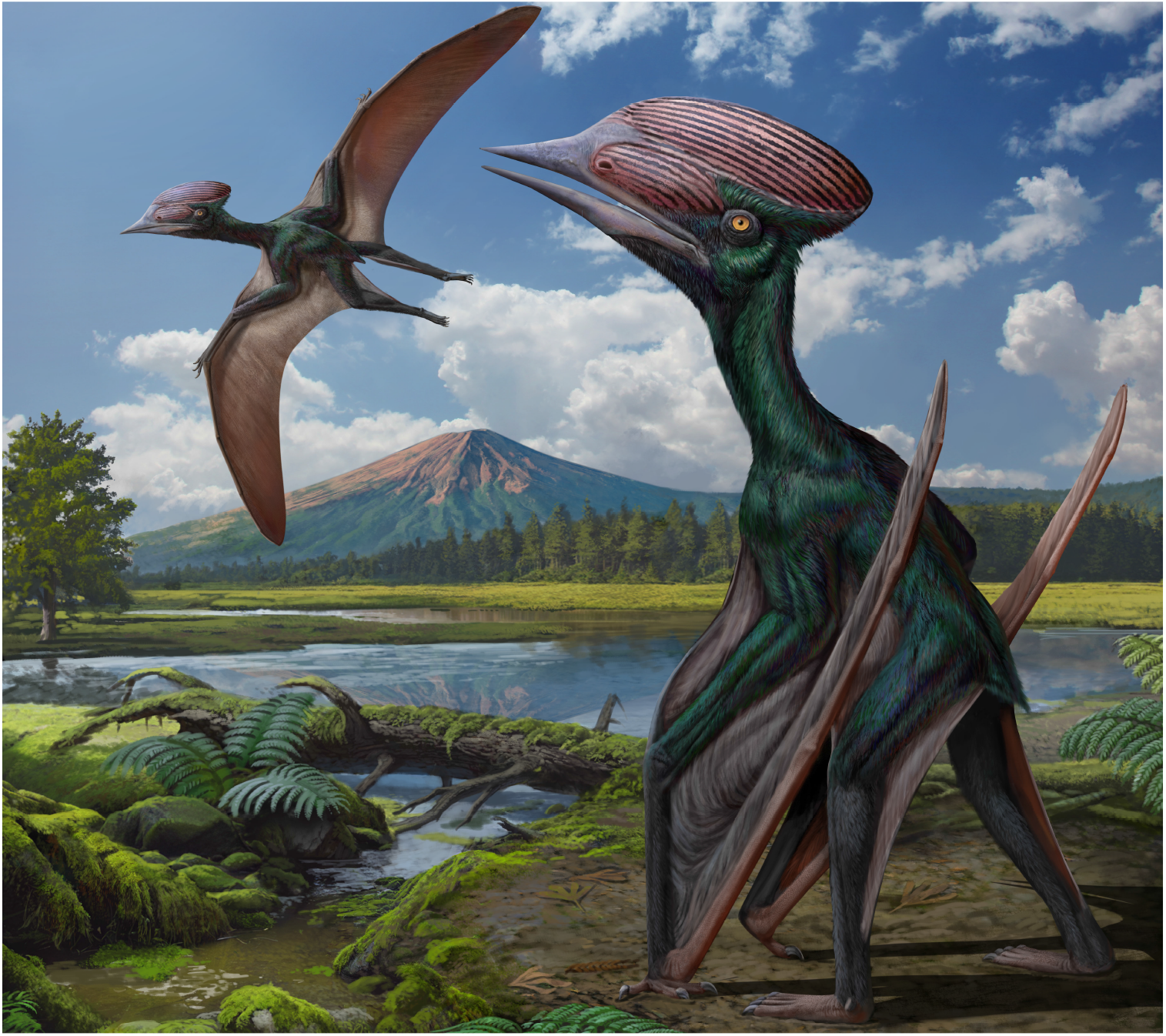
Reconstruction of the iridescent appearance of *Sinopterus dongi*. By courtesy of Chuang Zhao.

## Discussion

Monofilaments are widely distributed in theropods, ornithischian dinosaurs and pterosaurs (*20–25*). Whether these monofilaments in different taxa are homologous is crucial to understand the evolution of integumentary structures in Archosauria and the origin of feathers (*14, 15*). Our results document complex organization of melanosomes in pterosaur monofilaments (fig. S6, supplementary text) and a photonic structure similar to those seen in the barbs and barbules of extant avian feathers (Fig. 2).

Iridescent color has evolved independently multiple times in crown-group birds (*26, 27*), in part because melanosomes are likely organized through self-assembly processes (*28*). A hexagonal arrangement is the thermodynamically most stable position for tightly stacked rods (*29*) and thus a likely outcome of high melanosome density during development. Therefore, the large number of melanosomes below the photonic structure may have initiated self-assembly to form a highly ordered melanosome array (*28*). The size and morphology of the large melanosomes located in the center of monofilaments resemble the basal layer surrounding the medulla of extant non-iridescent structurally colored feathers in which color is formed by quasi-ordered keratin structure in the barb (L-3 in fig. S6, supplementary text) (*30*). But given the presence of the photonic structure and many disordered melanosomes below it, the specific function of these melanosomes should differ from that of the basal layer, which absorbs unscattered light. Indeed, the arrangement of melanosomes by size resembles those seen in hummingbirds that produce brilliant, highly iridescent colors (*31*). However, despite varying melanosome sizes, we see no evidence of the hollow platelet-like morphology of hummingbird melanosomes here.

Unlike the anurognathid and cf. *Jianchangnathus robustus* (*8, 12*), diverse melanosomes are reported only in tapejarid pterosaurs, including *Tupandactylus* cf. *imperator* (*13*) and *Sinopterus dongi* here. This indicates the regulatory mechanisms behind melanosome shape and organization, (e.g. melanocortin system and *PMEL17*) (*5, 6*), may have evolved new states in tapejarids. The previously reported low melanosome diversity in heterothermic organisms or those with low levels of homeothermy (*8*), contrasted with high diversity in high metabolic rate homeothermic organisms, i.e., birds and mammals, led to a suggested link between melanosome shape diversity and metabolic rate mediated through the melanocortin system (*8, 32*). While some pterosaurs showed low diversity (*8, 12*), others show high diversity (*13*). That tapejarid pterosaurs have diversity levels approaching those of birds and mammals is consistent with forms of homeothermy suggested for some early archosaurs (*33*). Indeed, pterosaur metabolism and that of early archosaurs have been repeatedly debated (*33, 34*). For example, both beta- and alpha-keratin monofilaments such as those in pterosaurs and mammaliaform taxa have been hypothesized to aid in insulation, and are not known in extant heterotherms. More independent data on thermoregulatory ability and melanosome diversity in pterosaurs is needed to further investigate these patterns.

Iridescent color in *Sinopterus dongi* provides new insight of the function of pycnofibres (Fig. 2,3). In extant birds, iridescent color feathers are highly correlated with intraspecific communication (*35*), especially courtship displays (*36*). Therefore, the iridescent monofilaments of *Sinopterus dongi* could also serve a visual signaling function. Sexually dimorphic cranial crests have been reported in pterosaurs e.g., *Pteranodon* and *Hamipterus* (*37, 38*), and the huge cranial crest of tapejarids was also hypothesized to have a display function (*39, 40*). The striated melanin-based pattern we found in the skin remnant near the premaxillary crest of *Sinopterus dongi* strongly supports this hypothesis (fig. S4). Sexual selection may play an important role in the evolution of pycnofibre coloration and cranial crest patterning in pterosaurs.

Previous reports of iridescent color in feathered dinosaurs and fossil birds based on melanosome morphology have led to the hypothesis that feathers functioned as sexual signals early in their evolution (*1, 9, 10, 26, 41–43*). However, these samples were limited to Paraves that already had complex feathers like extant birds. Unlike extant feathers, monofilaments are not involved in flight. Our results suggest that iridescent coloration may have been a part of the color repertoires of monofilaments that would have originated nearly 100 million years before the evolution of feathers, in the common ancestor of dinosaurs and pterosaurs. Alternatively, complex monofilaments, increases in melanosome diversity, and iridescent colors arose independently in Paraves and within Pterosauria from a flexible genetic suite common to Archosauria (*15*). The discovery of avian barb and barbule-style iridescent arrays in monofilaments suggests that a more complex gamut of coloration mechanisms may be available across Archosauria and deployed in sexual selection (*44*).

## Acknowledgments

We thank the Core Facilities of Life Sciences, Peking University for assistance with FIB-SEM and we would be grateful to J. Hu and Y.-Q. Liu for their help in sample testing; we thank D.-J. Tang and B.-Z. Xie at the FESEM Laboratory, China University of Geoscience, Beijing for their help in SEM and EDS analysis; C. Guo, Q. Li and Z.-P. Li (Tsinghua University) for their help in ToF-SIMS analysis; J. Yang (China University of Geoscience, Beijing) for XRF scanning of the specimens; C. Zhao for creating artistic reconstruction image; R. O. Prum for providing the Matlab code used in the two-dimensional fourier analysis.

## Funding

National Natural Science Foundation of China 42161134003 (Q.L., C.-F.Z., Z.W.)

National Natural Science Foundation of China 41872019 (Q.L., Z.W)

Fundamental Research Funds for the Central Universities for the Frontiers Science Center for Deep-time Digital Earth, China University of Geoscience (Beijing) 2652023001 (Q.L., Z.W.)

Chinese “111” project B20011 (Q.L.)

Taishan Scholar Program of Shandong Province tstp20240514 (C.-F.Z.)

Flemish Research Funds FWO G0E8322N and G0A7921N (M.D.S., L.D.)

Human Frontiers Science Program RGP0047 (M.D.S)

Air Force Office of Scientific Research FA9550-18-1-0477, FA9550-23-1-0622, and FA8655-23-2-7041 (M.D.S.)

## Author contributions

Conceptualization: L.D., Q.L., C.-F.Z., J.A.C., M.D.S.

Investigation: Z.W., J.L.

Visualization: Z.W., L.D.

Funding acquisition: L.D., C.-F.Z., Q.L., M.D.S.

Project administration: Q.L., Z.W.

Supervision: L.D., Q.L., C.-F.Z., M.D.S.

Writing – original draft: Z.W., L.D., M.D.S., Q.L.

Writing – review & editing: Z.W., L.D., C.-F.Z., J.A.C., J.L., M.D.S., Q.L.

## Data and materials availability

The fossil specimen CUGB-P2201 is accessioned at China University of Geoscience, Beijing, China (CUGB). All data are available in the main text, the supplementary materials, or DataDryad (https://datadryad.org/share/LINK_NOT_FOR_PUBLICATION/Y1v60b6QbXxG51ymYIaJaB_kIA_wjGByoXKtyyM6dAo).

## Supplementary Materials

Materials and Methods

Supplementary Text

Figs. S1 to S13

Tables S1 to S3

References (*45–97*)

Data S1

## References and Notes

1. Q. Li et al., Reconstruction of *Microraptor* and the evolution of iridescent plumage. Science 335, 1215–1219 (2012).

2. J. Vinther, D. E. G. Briggs, R. O. Prum, V. Saranathan, The colour of fossil feathers. Biol. Lett. 4, 522–525 (2008).

3. Q. Li et al., Plumage color patterns of an extinct dinosaur. Science 327, 1369–1372 (2010).

4. F. Zhang et al., Fossilized melanosomes and the colour of Cretaceous dinosaurs and birds. Nature 463, 1075–1078 (2010).

5. L. D’Alba, M. D. Shawkey, Melanosomes: biogenesis, properties, and evolution of an ancient organelle. Physiol. Rev. 99, 1–19 (2019).

6. M. E. McNamara et al., Decoding the evolution of melanin in vertebrates. Trends Ecol. Evol. 36, 430–443 (2021).

7. R. O. Prum, In Bird Coloration, Vol. 1, Mechanisms and measurements, G. E. Hill, K. J. McGraw, Ed. (Harvard Univ. Press, 2006), pp. 295–353.

8. Q. Li et al., Melanosome evolution indicates a key physiological shift within feathered dinosaurs. Nature 507, 350–353 (2014).

9. D. Hu et al., A bony-crested Jurassic dinosaur with evidence of iridescent plumage highlights complexity in early paravian evolution. Nat. Commun. 9, 217 (2018).

10. J. A. Peteya, J. A. Clarke, Q. Li, K. Q. Gao, M. D. Shawkey, The plumage and colouration of an enantiornithine bird from the Early Cretaceous of China. Palaeontology 60, 55–71 (2017).

11. A. W. A. Kellner et al. The soft tissue of *Jeholopterus* (Pterosauria, Anurognathidae, Batrachognathinae) and the structure of the pterosaur wing membrane. Proc. R. Soc. B: Biol. Sci. 277, 321–329 (2010).

12. Z. Yang et al., Pterosaur integumentary structures with complex feather-like branching. Nat. Ecol. Evol. 3, 24–30 (2019).

13. A. Cincotta et al., Pterosaur melanosomes support signalling functions for early feathers. Nature 604, 684–688 (2022).

14. J. A. Clarke, Feathers before flight. Science 340, 690–692 (2013).

15. C. B. Lowe, J. A. Clarke, A. J. Baker, D. Haussler, S. V. Edwards, Feather development genes and associated regulatory innovation predate the origin of Dinosauria. Mol. Biol. Evol. 32, 23–28 (2015).

16. S. A. Czerkas, Q. Ji, In Feathered Dinosaurs and the Origin of Flight, S. J. Czerkas, Ed. (The Dinosaur Museum, Blanding, 2002), pp. 15–41.

17. C. M. Eliason, M. D. Shawkey, A photonic heterostructure produces diverse iridescent colours in duck wing patches. J. R. Soc. Interface 9, 2279–2289 (2012).

18. C. Colleary et al., Chemical, experimental, and morphological evidence for diagenetically altered melanin in exceptionally preserved fossils. Proc. Natl. Acad. Sci. USA 112, 12592–12597 (2015).

19. M. E. McNamara, D. E. G. Briggs, P. J. Orr, D. J. Field, Z. Wang, Experimental maturation of feathers: implications for reconstructions of fossil feather colour. Biol. Lett. 9, 20130184 (2013).

20. M. J. Benton, D. Dhouailly, B. Jiang, M. E. McNamara, The early origin of feathers. Trends Ecol. Evol. 34, 856–869 (2019).

21. X. Xu et al., An integrative approach to understanding bird origins. Science 346, 1253293 (2014).

22. X. Xu, In The Evolution of Feathers, C. Foth, O. W. M. Rauhut, Ed. (Springer, 2020), pp. 67–78.

23. R. O. Prum, Development and evolutionary origin of feathers. J. Exp. Zool. 285, 291–306 (1999).

24. R. O. Prum, A. H. Brush, The evolutionary origin and diversification of feathers. Q. Rev. Biol. 77, 261–295 (2002).

25. C.-F. Chen et al., Development, regeneration, and evolution of feathers. Annu. Rev. Anim. Biosci. 3, 169–195 (2015).

26. K. K. Nordén et al., Melanosome diversity and convergence in the evolution of iridescent avian feathers-Implications for paleocolor reconstruction. Evolution 73, 15–27 (2019).

27. K. K. Nordén, C. M. Eliason, M. C. Stoddard, Evolution of brilliant iridescent feather nanostructures. Elife 10, e71179 (2021).

28. R. Maia, R. H. F. Macedo, M. D. Shawkey, Nanostructural self-assembly of iridescent feather barbules through depletion attraction of melanosomes during keratinization. J. R. Soc. Interface 9, 734–743 (2011).

29. J. Zhang, Y. Li, X. Zhang, B. Yang, Colloidal self-assembly meets nanofabrication: From two-dimensional colloidal crystals to nanostructure arrays. Adv. Mater. 22, 4249–4269 (2010).

30. F. Babarović et al., Characterization of melanosomes involved in the production of non-iridescent structural feather colours and their detection in the fossil record. J. R. Soc. Interface 16, 20180921 (2019).

31. M. A. Giraldo, J. L. Parra, D. G. Stavenga, Iridescent colouration of male Anna’s hummingbird (Calypte anna) caused by multilayered barbules. J. Comp. Physiol. A 204, 965–975 (2018)

32. C. M. Eliason, J. A. Clarke, Metabolic physiology explains macroevolutionary trends in the melanic colour system across amniotes. Proc. R. Soc. B: Biol. Sci. 285, 20182014 (2018).

33. M. J. Benton, The origin of endothermy in synapsids and archosaurs and arms races in the Triassic. Gondwana Res. 100, 261–289 (2021).

34. J. Wiemann et al., Fossil biomolecules reveal an avian metabolism in the ancestral dinosaur. Nature 606, 522–526 (2022).

35. S. M. Doucet, M. G. Meadows, Iridescence: a functional perspective. J. R. Soc. Interface 6, S115–S132 (2009).

36. J. Zi et al., Coloration strategies in peacock feathers. Proc. Natl. Acad. Sci. USA. 100, 12576–12578 (2003).

37. S. C. Bennett, Sexual dimorphism of Pteranodon and other pterosaurs, with comments on cranial crests. J. Vertebr. Paleontol. 12, 422–434 (1992).

38. X. Wang et al., Sexually dimorphic tridimensionally preserved pterosaurs and their eggs from China. Curr. Biol. 24, 1323–1330 (2014).

39. E. Frey, D. M. Martill, M.-C. Buchy, A new species of tapejarid pterosaur with soft-tissue head crest. Geol. Soc. Spec. Publ. 217, 65–72 (2003).

40. M. P. Witton, Pterosaurs Natural History, Evolution, Anatomy. (Princeton University Press, 2013).

41. C. M. Eliason, J. A. Clarke, Cassowary gloss and a novel form of structural color in birds. Sci. Adv. 6, eaba0187 (2020).

42. A. D. Croudace, C. Shen, J. Lü, S. L. Brusatte, J. Vinther, Iridescent plumage in a juvenile dromaeosaurid theropod dinosaur. Acta. Palaeontol. Pol. 68, 213–225 (2023).

43. Y. Pan et al., Unambiguous evidence of brilliant iridescent feather color from hollow melanosomes in an Early Cretaceous bird. Natl. Sci. Rev. 9, nwab227 (2022).

44. C. M. Eliason, J. A. Clarke, S. A. Kane, Wrinkle nanostructures generate a novel form of blue structural color in great argus flight feathers. Iscience 26, 105912 (2023).

45. Z. Yu, M. Wang, Y. Li, C. Deng, H. He, New geochronological constraints for the Lower Cretaceous Jiufotang Formation in Jianchang Basin, NE China, and their implications for the late Jehol Biota. Palaeogeogr. Palaeoclimatol. Palaeoecol. 583, 110657 (2021).

46. Ø. Hammer, D. A. T. Harper, P. D. Ryan, Past: paleontological statistics software package for education and data analysis. Palaeontol. Electronica 4, 1–9 (2001).

47. R Core Team, R: A Language and Environment for Statistical Computing, version 4.5.0, R Foundation for Statistical Computing, Vienna, Austria (2025); https://www.R-project.org/.

48. H. Wickham, J. Bryan, readxl, version 1.4.5, CRAN (2025); 10.32614/CRAN.package.readxl.

49. H. Wickham, D. Vaughan, M. Girlich, tidyr, version 1.3.1, CRAN (2024); 10.32614/CRAN.package.tidyr

50. H. Wickham et al., ggplot2, version 3.5.2, CRAN (2025); 10.32614/CRAN.package.ggplot2.

51. J. Ameijeiras-Alonso, R. M. Crujeiras, A. Rodríguez-Casal, multimode, version 1.5, CRAN (2021): 10.32614/CRAN.package.multimode.

52. L. Scrucca, C. Fraley, T. B. Murphy, A. E. Raftery, mclust, version 6.1.1, CRAN (2024); 10.32614/CRAN.package.mclust

53. R. O. Prum, R. H. Torres, A Fourier tool for the analysis of coherent light scattering by bio-optical nanostructures. Integr. Comp. Biol. 43, 591–602 (2003).

54. D. G. Stavenga, H. L. Leertouwer, D. C. Osorio, B. D. Wilts, High refractive index of melanin in shiny occipital feathers of a bird of paradise. Light. Sci. Appl. 4, e243–e243 (2015).

55. M. Xiao, A. Dhinojwala, M. D. Shawkey, Nanostructural basis of rainbow-like iridescence in common bronzewing Phaps chalcoptera feathers. Opt. Express 22, 14625–14636 (2014).

56. X. Wang, Z. Zhou, A new pterosaur (Pterodactyloidea, Tapejaridae) from the Early Cretaceous Jiufotang Formation of western Liaoning, China and its implications for biostratigraphy. Chin. Sci. Bull. 48, 16–23 (2003).

57. J. Lü, Y. Gao, L. Xing, Z. Li, Q. Ji, A new species of *Huaxiapterus* (Pterosauria: Tapejaridae) from the Early Cretaceous of western Liaoning, China. Acta Geol. Sinica Engl. Ed. 81, 683–687 (2007).

58. X. Zhang, S. Jiang, X. Cheng, X. Wang, New material of *Sinopterus* (Pterosauria, Tapejaridae) from the Early Cretaceous Jehol Biota of China. An. Acad. Bras. Ciênc. 91, e20180756 (2019).

59. R. V. Pêgas, X. Zhou, X. Jin, K. Wang, W. Ma, A taxonomic revision of the *Sinopterus* complex (Pterosauria, Tapejaridae) from the Early Cretaceous Jehol Biota, with the new genus Huaxiadraco. PeerJ 11, e14829 (2023).

60. X. Zhang et al., A new species of *Eopteranodon* (Pterodactyloidea, Tapejaridae) from the Lower Cretaceous Yixian Formation of China. Cretac. Res. 149, 105573 (2023).

61. E. Frey, H. Tischlinger, M.-C. Buchy, D. M. Martill, New specimens of Pterosauria (Reptilia) with soft parts with implications for pterosaurian anatomy and locomotion. Geol. Soc. Spec. Publ. 217, 233–266 (2003).

62. K. Eck, R. A. Elgin, E. Frey, On the osteology of *Tapejara wellnhoferi* Kellner 1989 and the first occurrence of a multiple specimen assemblage from the Santana Formation, Araripe Basin, NE-Brazil. Swiss J. Palaeontol. 130, 277–296 (2011).

63. J. Lü, C. Yuan, New tapejarid pterosaur from western Liaoning, China. Acta Geol. Sinica Engl. Ed. 79, 453–458 (2005).

64. J. Lü et al., A new species of *Huaxiapterus* (Pterosauria: Pterodactyloidea) from the Lower Cretaceous of western Liaoning, China with comments on the systematics of tapejarid pterosaurs. Acta Geol. Sinica Engl. Ed. 80, 315–326 (2006).

65. J. Lü et al., New material of pterosaur *Sinopterus* (Reptilia: Pterosauria) from the Early Cretaceous Jiufotang Formation, western Liaoning, China. Acta Geol. Sinica Engl. Ed. 80, 783–789 (2006).

66. C.-F. Zhou, T. Niu, D. Yu, New data on the postcranial skeleton of the tapejarid *Sinopterus* from the Early Cretaceous Jehol Biota. Hist. Biol. 35, 356–363 (2022).

67. C.-F. Zhou, C. Miao, B. Andres, New data on the cranial morphology of the tapejarid *Sinopterus* from the Early Cretaceous Jehol Biota. Hist. Biol. 36, 1–8 (2023).

68. J. Lü et al., The toothless pterosaurs from China. Acta Geol. Sinica 90, 2513–2525 (2016).

69. C.-F. Zhou, D. Yu, Z. Zhu, B. Andres, A new wing skeleton of the Jehol tapejarid Sinopterus and its implications for ontogeny and paleoecology of the Tapejaridae. Sci. Rep. 12, 10159 (2022).

70. V. Beccari et al., Osteology of an exceptionally well-preserved tapejarid skeleton from Brazil: Revealing the anatomy of a curious pterodactyloid clade. PloS one 16, e0254789 (2021).

71. A. W. A. Kellner, Pterosaur phylogeny and comments on the evolutionary history of the group. Geol. Soc. Spec. Publ. 217, 105–137 (2003).

72. D. M. Unwin, On the phylogeny and evolutionary history of pterosaurs. Geol. Soc. Spec. Publ. 217, 139–190 (2003).

73. A. W. A. Kellner, New information on the Tapejaridae (Pterosauria, Pterodactyloidea) and discussion of the relationships of this clade. Ameghiniana 41, 521–534 (2004).

74. P. C. Manzig et al., Discovery of a rare pterosaur bone bed in a Cretaceous desert with insights on ontogeny and behavior of flying reptiles. PloS one 9, e100005 (2014).

75. A. W. A. Kellner, L. C. Weinschütz, B. Holgado, R. A. M. Bantim, J. M. Sayão, A new toothless pterosaur (Pterodactyloidea) from southern Brazil with insights into the paleoecology of a Cretaceous desert. An. Acad. Bras. Ciênc. 91, e20190768 (2019).

76. J. J. Li, J. C. Lü, B. K. Zhang, A new Lower Cretaceous sinopterid pterosaur from the western Liaoning, China. Acta Palaeontol. Sinica 42, 442–447 (2003).

77. J. Lü, B. Zhang, New pterodactyloid pterosaur from the Yixian Formation of western Liaoning. Geol. Rev. 51, 458–462 (2005).

78. X. Wang, Z. Zhou, Pterosaur assemblages of the Jehol Biota and their implication for the Early Cretaceous pterosaur radiation. Geolog. J. 41, 405–418 (2006).

79. D. Naish, M. P. Witton, E. Martin-Silverstone, Powered flight in hatchling pterosaurs: evidence from wing form and bone strength. Sci. Rep. 11, 13130 (2021).

80. S. C. Bennett, The ontogeny of Pteranodon and other pterosaurs. Paleobiology 19, 92–106 (1993).

81. A. W. A. Kellner, Comments on Triassic pterosaurs with discussion about ontogeny and description of new taxa. An. Acad. Bras. Ciênc. 87, 669–689 (2015).

82. J. A. Wilson, Anatomical nomenclature of fossil vertebrates: standardized terms or ‘lingua franca’? J. Vertebr. Paleontol. 26, 511–518 (2006).

83. S. C. Bennett, The osteology and functional morphology of the Late Cretaceous pterosaur *Pteranodon* Part I. General description of osteology. Palaeontogr. Abt. A. 260, 1–112 (2001).

84. L. P. A. M. Claessens, P. M. O’Connor, D. M. Unwin, Respiratory evolution facilitated the origin of pterosaur flight and aerial gigantism. PloS one 4, e4497 (2009).

85. X. Wang, Z. Zhou, F. Zhang, X. Xu, A nearly completely articulated rhamphorhynchoid pterosaur with exceptionally well-preserved wing membranes and “hairs” from Inner Mongolia, Northeast China. Chin. Sci. Bull. 47, 226–230 (2002).

86. J. C. Lü, Soft tissue in an Early Cretaceous pterosaur from Lianoning Province, China. Memoir of the Fukui Prefectural Dinosaur Museum 1, 19–28 (2002).

87. D. M. Unwin, N. N. Bakhurina, *Sordes pilosus* and the nature of the pterosaur flight apparatus. Nature 371, 62–64 (1994).

88. X. Xu, Z. L. Tang, X. L. Wang, A therizinosauroid dinosaur with integumentary structures from China. Nature 399, 350–354 (1999).

89. X. T. Zheng, H. L. You, X. Xu, Z. M. Dong, An Early Cretaceous heterodontosaurid dinosaur with filamentous integumentary structures. Nature 458, 333–336 (2009).

90. J. Lindgren et al., Molecular preservation of the pigment melanin in fossil melanosomes. Nat. Commun. 3, 824 (2012).

91. J. Lindgren et al., Molecular composition and ultrastructure of Jurassic paravian feathers. Sci. Rep. 5, 13520 (2015).

92. A. Roy et al., In Pennaraptoran theropod dinosaurs: past progress and new frontiers, M. Pittman, X. Xu, Ed. (American Museum of Natural History, New York, 2020), pp. 251–276.

93. L. J. Preston et al., The preservation and degradation of filamentous bacteria and biomolecules within iron oxide deposits at Rio Tinto, Spain. Geobiology 9, 233–249 (2011).

94. M. E. McNamara et al., Fossilized skin reveals coevolution with feathers and metabolism in feathered dinosaurs and early birds. Nat. Commun. 9, 2072 (2018).

95. Z. Yang, B. Jiang, J. Xu, M. E. McNamara, Cellular structure of dinosaur scales reveals retention of reptile-type skin during the evolutionary transition to feathers. Nat. Commun. 15, 4063 (2024).

96. J. A. Clarke et al., Fossil evidence for evolution of the shape and color of penguin feathers. Science 330, 954–957 (2010).

97. J. Vinther, A guide to the field of palaeo colour: Melanin and other pigments can fossilise: Reconstructing colour patterns from ancient organisms can give new insights to ecology and behaviour. Bioessays 37, 643–656 (2015).

