## supplemental files for "Iridescence in pterosaur pycnofibers and the evolution of integumentary coloration"

### Materials and Methods

#### Specimen provenance and sampling

The specimen CUGB-P2201 is composed of four slabs with bones and soft tissue. This specimen was collected by author Quanguo Li in 2022 from the Lower Cretaceous Jiufotang Formation at Lamadong locality, Jianchang county, Huludao city, Liaoning Province, China. The zircon U-Pb dating of the outcrops in this area is about 118.9 Ma (*45*), which corresponds to the Aptian, Early Cretaceous. Pictures of this specimen were taken with a Nikon D800 camera, and close-ups of soft tissue were taken with a Zeiss SteREO Discovery. V20 stereo microscope. The length of bones was obtained by measuring Sr distribution image of the micro-XRF with software ImageJ (available for download at https://imagej.net/ij/) and verified by measuring the specimen directly using vernier calipers (table S1). See supplementary information for a detailed description of this specimen. We collected small samples (less than 2.5 $mm^2$) from different areas of soft tissue, including head crest soft tissue and monofilaments. During these sampling we selected separated regions to prevent potential influence of skin or internal tissues.

#### SEM and EDS analysis

We performed scanning electron microscopy (SEM) to observe the microstructure of soft tissues. Prior to SEM experiment, the samples were fixed on double-sides carbon tape and Au/Pt sputter-coated (approximately 10 nm thickness) in a Leica EM ACE600 sputter coater, these samples were examined using a ZEISS SUPRA-55VP field emission scanning electron microscope at the FESEM Laboratory, China University of Geoscience, Beijing, with a working distance of 8.7-20 mm and accelerating voltage of 10-20 kV. Images were acquired in secondary electron or backscatter mode. The elemental composition of selected micron-sized spots was analyzed by an Oxford X-act energy dispersive X-ray spectrometer (EDS) connected to the FESEM with accelerating voltage of 20 kV. The elemental distribution of these samples was analyzed by an Oxford Ultim Max 170 energy dispersive X-ray spectrometer connected to a Zeiss Crossbeam 550 focused ion beam scanning electron microscope at the Key Laboratory of Deep Petroleum Intelligent Exploration and Development, Institute of Geology and Geophysics, Chinese Academy of Sciences.

#### FIB-SEM

To explore the internal structure of monofilaments, we performed dual-beam focused ion-beam scanning electron microscopy (FIB-SEM) experiment. The expereiments were performed using an FEI Helios Nanolab G3 UC dual-beam FIB-SEM at the Core Facilities of Life Sciences, Peking University (Beijing, China), before the experiments, the samples were Au/Pt sputter-coated in a Harachi MC1000 sputter coater for 120 s. We selected well-preserved regions in monofilaments under scanning electron microscope and subsequently cut longitudinal sections as well as cross sections using the ion beam. Different beam conditions were chosen for different sized regions, initial FIB-SEM trench cuts were usually performed with a gallium-ion beam at accelerating voltage of 30 kV and current of 0.43/2.5/9.3 nA, followed by repeated cuts of the sections under the same conditions or with reduced current to eliminate curtaining/waterfall artifacts. The accelerating voltage for final imaging was 2 kV and the current was 0.2 nA. Since there is a 52° angle between the electron beam and the ion beam of the FIB-SEM, which will cause longitudinal deformation during imaging due to tilted viewing angles, we performed tilt-correction during imaging, or used the image-processing program ImageJ to adjust the height of the images to 1.269 times (1/sin 52°) to achieve the same correction effect. The image resolution was ensured using bicubic interpolation during the adjustment.

**Measurement of melanosomes, photonic structure and monofilament thickness**

We used the image-processing program ImageJ to accomplish the measurement of melanosomes morphology. We measured the maximum long axis and short axis length of melanosomes in the direction perpendicular to the FIB-SEM longitudinal sections and calculated the aspect ratio (long axis/short axis). When measuring cross sections, we selected melanosomes that were perpendicular to the cross section (i.e., individuals with a morphology close to circle) and measured their short axis length to obtain accurate diameter data. For statistics, we calculated the mean and standard deviation of the melanosome morphology data in different laminae, scatter plot and box plots of the data were plotted using the freeware PAST (*46*) (version 4.11, available for download at https://www.nhm.uio.no/english/research/resources/past/). Data on melanosomes of extant avian feathers were obtained from Li et al., 2012 (*1*), Babarović et al., 2019 (*30*) and Nordén et al., 2019 (*26*) (we selected only cylindrical melanosomes from iridescent feathers because flat melanosomes were not found in this specimen). To measure the parameters of photonic structure, we selected cross sections at M2csls4 corresponding to Fig. 2F (including 27 images across 8 sections from M2cs4(1).s1-M2cs4(1).s8). Measurements were performed on regions within cross sections meeting the following criteria: (1) the distance between the region and the overlapping boundary (i.e., the violet dashed line labeled in Fig. 2F,I) was within 15 melanosomes (since Fig. 2C,F both show approximately 15 layers of ordered melanosomes, measurements were confined to the corresponding regions); (2) Melanosomes exhibit circular or near-circular shapes (ensuring parallel alignment within the region and perpendicularity to cross section); (3) Melanosomes are densely packed with no significant gaps between them. We measured the distance between the centers of the neighboring melanosomes as the lattice spacing. For the potential "surface" layer of monofilament (fig. S8Q-S, U-X, see below), we measured the thickness of "surface" layer at position M17ls1(fig. S8Q-S) and M24cs1 (fig. S8W-X) as a criterion for the potential surface on the dorsal and ventral sides of the monofilament. When estimating the morphology of the monofilaments, we counted the maximum thickness of different melanosomes laminae preserved in sections to estimate the thickness of an intact monofilament. Since the sections were not cut perpendicular to the monofilament surface, we corrected for the thickness of different melanosomes laminae and potential "surface" layer: the actual thickness = measured thickness × cos$\theta$ ($\theta$ is the angle between the section and the vertical plane of the monofilament surface, 0°, 32° or 52° in this paper).

**Statistical analysis**

To investigate the internal structure and distribution of melanosomes in monofilaments, we performed multimodality testing and Gaussian mixture model fitting on melanosome diameter data collected from cross-sections of position M2csls1 and sections of position M16ls1. Analyses were performed in R (v4.5.0) (*47*), data processing and import utilized the R packages "readxl" (v1.4.5) (*48*) and "tidyr" (v1.3.1) (49), figures were generated with "ggplot2" (v3.5.2) (50). Multimodality test was performed using R package "multimode" (v1.5) (51), employing the "ACR" method. For unimodality testing, the replicate number "B" was set to 2000; for bimodality testing, "B" was set to 1000. Gaussian mixture model fitting was implemented using the R package "mclust" (v6.1.1) (52), with the "Modelnames" set to V (unequal variance).

**Two-dimensional Fourier analysis**

To further demonstrate the photonic structure in monofilaments, we used an established two-dimensional Fourier analysis tool (*53*) to characterize the periodicity of melanosomes arrangement in different directions. We converted FIB-SEM section images to grayscale mode before analysis and then analyzed them using MATLAB (version 9.10.0) and the 2-D Fourier tool (*53*). (In this tool, the average brightness of the image is subtracted to eliminate the effect of the central peak). The refractive index of melanin and keratin in this analysis were set to 1.74 and 1.54, respectively (*54*).

**Color simulation**

We built three-dimensional models using dimensions extracted from electron micrographs in a commercial Maxwell equation solver from ANSYS Solutions, Inc. varied the angle of incident light for each model from 90° to 40°, predicting that this would cause variation in the predicted reflectance curves. We applied PML boundary conditions for the X-axis and periodic boundary conditions along the Y- and Z- axis to obtain all the light reflected by the structure, a polarized plane wave light source with normal incidence, and acquired the reflectance spectra for the visible wavelengths (400-700 nm).

We created seven models of melanosome laminae inside monofilaments. Model 1 is a 'simulation cell' containing 15 layers of hexagonal melanosome photonic crystals (length = 606.5 nm, diameter = 147 nm, lattice spacing = 187 nm) in a keratin matrix (Fig. 3C). Model 2 adds a potential surface keratin cortex to Model 1 (thickness = 381.7 nm) to represent the "surface" observed in some monofilament locations (Fig. 3A). For Model 3 we repeated Model 2 but increased the size of melanosomes by 10% (fig. S13A), accounting for the potential taphonomic shrinkage of melanosomes during diagenetic process. We added disordered melanosomes and a uniform melanin layer at the bottom of the melanosome photonic crystals in Model 1-Model 3 to simulate the effect of melanosomes inside the monofilament on the reflectance spectrum. Additionally, we modelled the melanosome array directly from FIB-SEM images in a few locations within the monofilaments, including Model 4 (with and without the cortex, Fig. 3B and Fig. 3D, based on Fig. 2I), Model 5 (without the cortex, fig. S13B, based on fig. S10D), Model 6 (without the cortex, fig. S13C, based on fig. S10E) and Model 7 (without the cortex, fig. S13D, based on fig. S10F). We used perfectly matched layers to absorb electromagnetic waves from the top and bottom of the simulation cell and periodic boundary conditions on the sides to simulate infinite periodic structures. We used previously reported values of the refractive index of keratin (*55*) and the wavelength-dependent RI and absorption of melanin (*55*). The light source covered a wavelength range of 300-700 nm and corresponded to an unpolarized plane wave at normal incidence unless otherwise indicated.

**Micro-XRF**

The specimen CUGB-P2201 was scanned using a Bruker M6 Jetstream micro X-ray fluorescence spectrometer (micro-XRF) at China University of Geoscience, Beijing. This instrument allows the analysis of micro-zones on large samples and is therefore suitable for fossils. The M6 Jetstream instrument consists of a mobile measuring head and a XY-motorized stage, the measuring head is equipped with a Rh-target X-ray tube (typically operating at 50 kV and 600 μA) and the X-ray beam is guided through a polycapillary focusing optics to the sample surface with a spot size of 50 μm. The final signal is captured by a silicon drift detector, this detector is 50 $mm^2$ with energy resolution < 145 eV at Mn-Kα. Due to the fact that specimen CUGB-P2201 are preserved scattered and at a distance from each other, our scans were carried out in several passes. The data acquisition speed was 10 ms/pixel and the resolution was set to 100 μm. The sizes of different scanning regions were 4779 × 2581 pixels (fig. S3Aa), 3294 × 1887 pixels (fig. S3Ab), 1883 × 700 pixels (fig. S3Ad), 1227 × 370 pixels (fig. S3Ac) respectively, and the total scanning time was 47 h 58 min. After scanning, we used the companion software Esprit M6 to analyze the XRF spectra and generate the element distribution maps as well as heat maps. The color shades of the element distribution maps indicate the degree of elemental enrichment, and heat maps show the enrichment of single element through color tones.

**ToF- SIMS**

The ToF-SIMS analysis was done at Tsinghua University with a ToF-SIMS 5-100 instrument (ION-TOF GmbH, Germany) equipped with a Bi liquid metal ion gun (LMIG) to investigate the chemical composition of monofilaments and skin remnant near the premaxillary. ToF-SIMS spectra were acquired using 30keV $Bi_3^{++}$ primary ions and low energy electron flooding for charge compensation. The scanning area was set to 200 × 200 μm$^2$ and 50 × 50 μm$^2$ to investigate the distribution of eumelanin related ions in relation to soft tissue and the ToF-SIMS spectra excluding the effect of sediment. The surface of the skin remnant region has sediments, and the melanosomes are usually not directly exposed, so we used GCIB (Gas Cluster Ion Beam) to sputter the surface before analyzing it, sputtering rate was 0.13 nm/s for $SiO_2$.

**FIB-ToF-SIMS**

We conducted FIB-ToF-SIMS analysis at the Key Laboratory of Deep Petroleum Intelligent Exploration and Development, Institute of Geology and Geophysics, Chinese Academy of Sciences to probe the chemical composition of potential "surface" found in monofilaments (see supplementary text for description of monofilament surfaces). The instrument is a ZEISS Crossbeam 550 focused ion beam scanning electron microscope equipped with a retractable ToF-SIMS spectrometer. This instrument can accurately localize the analyzed region under SEM and can indicate the distribution of secondary ions at different depth, thus providing accurate information about the monofilament surface. ToF-SIMS spectra were acquired using 30keV, 50pA Ga primary ions. The analytical areas of positive and negative ion spectras are 3.8 × 3.8 μm$^2$ and 4.2 × 4.2 μm$^2$, respectively.

### Supplementary Text

#### Anatomical description of CUGB-P2201

The specimen CUGB-P2201 is a partially disarticulated skeleton, consisting of the cranial crest, some vertebrae and most of the limb bones. The skeleton is heavily crushed as in two-dimensional preservation. Soft tissues are distributed around the skeleton.

***Premaxilla***

The premaxillary crest is partially preserved in right view with a length of approximately 166.7 mm (fig. S1E). It has a slightly concave dorsal margin and a pointed posterior end. The upturned morphology of the posterior crest is similar to the other Jehol tapejarids (*56–60*). Along the dorsal margin of the premaxillary crest, many fine crest fibres are well preserved (fig. S1E, I-K). These fibres are tightly packed with roughly 8 fibres / mm and the longest one exceeds 5.81 mm. At the anterior part of the crest, the fibres are nearly perpendicular to the bony crest. Posteriorly, the fibres are gradually inclined, and almost parallel to the bony crest at the posterior end (fig. S1I-K). The fibres are partially ossified proximally, similar to previously reported Brazilian tapejarid *Tupandactylus navigans* and *T. imperator* (*61*).

***Cervical vertebra***

Only one cervical vertebra is preserved and exposed in dorsal view, which belongs to the middle cervical series (fig. S2Q,R). It is approximately 42.37 mm long (from tip of prezygapophysis to the distal margin of centrum) and 16.63 mm wide (at the centrum most constricted part), with an aspect ratio of about 2.5. The left prezygapophysis is incomplete whereas the right is well-preserved, it is horny-like, forms an angle of about 25° with the sagittal plane in dorsal view. The postzygapophysis projects more laterally and has a relatively higher position than the prezygapophysis. In anterior view, there is a pneumatic foramen adjacent to the right side of the neural canal, whereas the other side is obscure by the compression. The lateral pneumatic foramen is unrecognizable because of compression.

***Dorsal vertebrae***

Three articulated dorsal vertebrae, two isolated neural arches and three isolated centra are preserved (Fig. 1A,B). The articulated dorsal vertebrae are exposed in dorsal view (Fig. 1A,B), two of which are structurally intact, the third one only preserves a centrum. The transverse process of the anterior two dorsal vertebrae protrudes laterally at a nearly perpendicular angle. The complete isolated neural arch is preserved near the distal end of wing metacarpal and exposed in anterior view (fig. S2S,T). The diameter of the neural canal is nearly 2 mm. The slender transverse process protrudes dorsolaterally. The neural spine is nearly 8 mm high, and widens ventrally, forming a triangular profile in anterior view. A large pneumatic foramen is located at the base of the neural spine, and a small foramen is present on the right of the base. Three procoelous centra show a similar morphology with an aspect ratio of about 1.2 (Fig. 1A,B, fig. S2U,V). The two shallow ridges of these centra that frame the neural canal are clearly visible. The separation of the neural arch and centrum indicates they are not fused, which is a feature of immaturity.

***Sternum***

The sternum is incompletely preserved, and exposed in dorsal view (Fig. 1A,B). The preserved part consists of a sternal plate and a broken cristospine. The sternal plate is slightly concave.

***Pectoral girdle***

The scapula and coracoid are scattered (Fig. 1A,B). The left scapula is a blade-like bone exposed in ventral view and broken into two pieces. The proximal part of the left scapula expands to form an articular facet for the coracoid. It curves medially and forms an angle of about 138° with the shaft. A distinct depression is observed in the proximal center of the scapula, but no pneumatic foramen can be observed. The scapular shaft is straight with constant width about 9.5 mm. The right scapula is preserved near the left one and also exposed in ventral view. Its preserved length is 62.2 mm. The distal scapular shaft has a constant width of about 11.5 mm, and the angle between the proximal bend and shaft is approximately 150°.

The left coracoid is complete and exposed in dorsal view (fig. S2K,L). Its total length is 55.5 mm. The shaft is constricted in the middle position. The distal end expands and forms a "saddle-like" articulation for the sternum. The proximal end of the coracoid expands to form an articulation facet for the scapula. Ventrally, a pneumatic foramen is positioned on the coracoid glenoid cavity. The proximal expansion of the coracoid exceeds that of *Tapejara wellnhoferi* (*62*), and more closely resembles the condition seen in other Jehol tapejarids (*56*, *59*, *63–67*).

***Humerus***

Both humeri are preserved in ventral view. The left humerus is incomplete with its ulnar crest and humeral head not preserved (fig. S2A,B). The deltopectoral crest is well-developed. It is straight, unwarped and perpendicular to the humeral shaft. The extended length of the deltopectoral crest is 19.9 mm with a base of 18.9 mm wide. The anterior margin of the deltopectoral crest is slightly rounded. The proximal and distal margins are parallel to each other, which leads to a subrectangular profile. The humeral shaft is long and straight with a slightly expanded distal end. Two separated epiphyses are preserved against the humerus, suggesting this specimen was in a juvenile stage. The larger epiphyses is nearly elliptical with a concave center, while the smaller one is nearly spherical. The right humerus is completely preserved (fig. S2C,D), but obscured by the right ulna and the left first wing phalanx. Its length is 83.42 mm. The subrectangular deltopectoral crest is about 18.0 mm long, and 17.2 mm wide at its base. The humeral head is subtriangular with a proximal tip obscured by the carpals. A large ventral pneumatic foramen is present near the head, resembles the other tapejarids (*62–63*, *68–70*). The humeral shaft is straight with a diameter of about 13 mm. The distal condyles are separated by an intercondylar sulcus.

***Radius and ulna***

The left ulna and radius are exposed in ventral view with the distal ends missing (Fig. 1A,B, fig. S2A,B). The left ulna is estimated to be 130.2 mm long based on its preserved impressions. It expands proximally and slightly retracts toward the distal end. The left radius is parallel to the ulna and obscured proximally by the ulna and humerus. The estimated length of left radius is about 125.28 mm. The diameter of the ulna is larger than the radius.

***Carpals***

The proximal and distal carpals of the left forelimb are preserved in situ at the distal end of the radius and ulna (Fig. 1A,B). It is impossible to determine whether the proximal / distal carpals have fused into proximal / distal syncarpal due to the overlapping of the first wing phalanx, but the suture between the proximal and distal carpals is clear. The right pteroid is rod-like and has a length of 65.8 mm, nearly 50% of the ulna (Fig. 1A,B).

***Metacarpals***

The left metacarpal I-III are rod-shaped and slightly expanded at the distal end to articulate the manual digits (Fig. 1A,B), their exact length and contact with carpals are unavailable.

The metacarpal IV (wing metacarpal) is significantly larger than the first three. The left metacarpal IV is exposed in posterior view. It is long and straight with an estimated length of 141.06 mm. The metacarpal IV is approximately 1.08 times length of the left ulna, with a ratio of 0.81 to the first wing phalanx. The metacarpal IV bears a proximal expansion and gradually contracts distally. The distal end is slightly enlarged to form two subequal condyles. The ventral condyle is almost parallel to the metacarpal IV, whereas the dorsal condyle is expanded dorsally. There is a large pneumatic foramen positioned between two condyles, as in *Tapejara wellnhofer* (*62*) and other Jehol tapejarids (*66*, *69*). The right metacarpal IV is incompletely preserved.

***Manual digit I-III***

Only the left manual digits I-III are preserved, showing the typical phalangeal formula of "2-3-4" (fig. S2G,H). The length is gradually increasing from digit I to III. The first phalanx of digit III is the most robust element. The second phalanx of digit III is a short rectangular element, with an aspect ratio of about 1.5. The remaining phalanges are rod-shaped and slightly expand at the proximal and distal ends to form articular facets. The manual unguals (the ultimate phalanges) are large and more sharply curved than the pedal unguals. The curvature is approximately 97° along the inner margin. Proximally, the unguals are expanded with a prominent flexor tubercle occupying almost half of the proximal end. A distinct midline groove develops along the distal margin of the unguals. Black keratinous sheath is preserved at the distal end of the second and third unguals, extending more than half the length of the bony element (fig. S2I,J).

***Wing finger (manual digit IV)***

The wing phalanges are incompletely preserved. The left first wing phalanx is a straight and rod-shaped bone, exposed in dorsal view (Fig. 1A,B). The proximal end of the first wing phalanx expands anteroposteriorly to form an articular facet for the wing metacarpal. The extensor tendon process is unfused with the first wing phalanx, suggesting this individual is still immature. The distal end of this phalanx is posteriorly expanded relative to shaft to form the broad articular facet for the second wing phalanx. The left second wing phalanx is broken into three pieces and exposed dorsally. Its proximal end is expanded, the shaft is straight and with constant thickness. The left third wing phalanx is preserved with only the proximal end. The fourth wing phalanx is not preserved.

The right-wing finger is quite fragmentary, detailed information from the first wing phalanx to the second wing phalanx is not recognizable. The articulation between the right third wing phalanx and fourth wing phalanx is clear. The fourth wing phalanx is about 46.6 mm long, with a ratio of 0.27 to the left first wing phalanx. This proportion aligns with *Sinopterus dongi* and *Huaxiapterus jii* (*56*, *63*), but is significantly larger than that of *Huaxiapterus corollatus* and *Huaxiapterus benxiensis* (*57*, *64*).

***Pelvic girdle***

The left pubis and right ischium are preserved. The left pubis is preserved in lateral view (fig. S2M,N). It is subquadrilateral with a concave anterior margin, slightly convex ventral margin, and nearly straight posterior margin. The pubis is widened along the acetabulum, resulting in the dorsal margin clearly exceeding the pubic plate. The strongly concave anterior margin is different from that of Brazil tapejarids *Tapejara wellnhofer* (*62*), but similar to the other Jehol tapejarid *Sinopterus* (*59*, *66*). A medium-sized obturator foramen is located on the posterior margin of the pubis.

The right ischium is exposed in lateral view with a subtriangular shape (fig. S2O,P). Its surface is flat and smooth. The ischium is convex dorsally and elongated anteroposteriorly, reaching its longest part at the nearly straight ventral margin. The posterior margin is curved with a distinctly convex posteroventral corner. The anteroventral part retains the original contour impression. A small tuberculum is located at the anterodorsally one third position of this bone.

***Femur***

Both femora are preserved. The left femur is completely preserved and exposed in medial view (Fig. 1A,B). The femoral head is semi-spherical and tilted relative to the neck. The femoral shaft is bowed anteriorly. The distal end of the femur contains two condyles in which the lateral condyle is significantly larger than the medial one.

The right femur is exposed in posterior view. Its proximal and distal portions are well preserved. The shaft is severely crushed (Fig. 1A,B), creating the illusion that the right femur looks longer than the left. The proximal part of the right femur preserves the femoral head and neck, which forms an angle of 140° with the shaft. On the lateral side of the femoral neck, the greater trochanter is well developed, but no pneumatic foramen exists between the femoral neck and the greater trochanter. The distal end of the right femur is expanded to form an articular facet for the tibia, which consists of a pair of condyles and a broad intercondylar sulcus. The lateral condyle is incompletely preserved, while the medial one is intact.

***Tibia and fibula***

Both sides of the tibia and fibula are incompletely preserved (Fig. 1A,B). The left tibia and fibula are still articulated with the femur, only the proximal part of which are preserved and exposed in posterior view. The tibia is a long and rod-like bone whose articulation with femur is quite wide. The left fibula is preserved parallel to the tibia, its proximal end reaches the articular surface of tibia.

Both the proximal and distal ends of the right tibia are incomplete. The shaft is straight. The right fibula is partially preserved and aligned parallel to the tibia.

***Tarsus***

One proximal tarsal and two distal tarsals of the right pes are preserved (fig. S2E,F). The proximal tarsal is not fused with the tibia, suggesting this is an immature individual. The proximal tarsal is distinctly larger than the two distal tarsals with a subrectangular shape. The lateral distal tarsal is subellipsoidal with a smooth margin and located distally against the metatarsal IV. The medial one has an irregular shape and is preserved against metatarsal II-III distally.

***Metatarsus***

The right metatarsals are well preserved (fig. S2E,F). The metatarsals I-IV are long, slender and parallel to each other. The metatarsals I and II are almost equal in length, whereas the metatarsals III and IV are progressively shorter in length. It is different in diameter of the metatarsals, although the diameters of metatarsals I – IV are nearly the same at the distal part. Metatarsal I and metatarsal IV expanded proximally, whereas metatarsal II and metatarsal III do not show any change proximally in diameter. The metatarsal V is stout and sub-triangular in shape. It is strongly shortened with a total length of only 9.4 mm, equivalent to 26.7% of the metatarsal I.

***Pedal digits***

The pedal digits are completely preserved with a phalangeal formula of 2-3-4-5-1 (fig. S2E,F). Among digits I-IV, the second phalanx of digit III, the second and the third phalanges of digit IV are dramatically shortened, while the remaining phalanges are rod-like, with thickening proximal and distal ends to form the articular facet. The digit V degenerates into a single nubbin-like phalanx, a synapomorphy of the pterodactyloids. The pedal unguals are smaller in size and curved less than the manual unguals. The curvature of pedal unguals is approximately 56° along the inner margin. At the base of pedal unguals, the flexor tubercle is weakly developed. There are obvious midline-grooves present on the lateral surfaces of the distal unguals, similar to the manual unguals.

**Taxonomy**

This new specimen exhibits some typical features of Pterodactyloidea, including elongated wing metacarpal, highly developed humeral deltopectoral crest and degenerated pedal digit V (*71*, *72*). Furthermore, the subrectangular and unwarped deltopectoral crest shape, moderately elongated mid-cervical, and the well-developed cranial sagittal crest support this specimen as a taxon of Tapejaridae (*71*, *73*). On this basis, the enlarged-proximally coracoid, the upturned distal end of the premaxillary crest, the shape of pubis and the proportions of limb bones differ considerably from Brazil tapejarids (*62*, *70*, *74*, *75*), but similar to the Jehol tapejarids (*56*, *58*, *67*).

Hitherto, a total of nine tapejarid species have been named from the Jehol Biota. Seven of these species are reported from the Jiufotang Formation: *Sinopterus dongi* (*56*), *S. gui* (*76*), *S. lingyuanensis* (*68*), *Huaxiapterus jii* (*63*), *H. corollatus* (*64*), *H. benxiensis* (*57*) and *H. atavismus* (*68*), and two species have been named from the Yixian Formation: *Eopteranodon lii* (*77*) and *E. yixianensis* (*60*). There is still controversy over the classification of these Jehol tapejarids (*39*, *58*, *59*, *78–79*). Based on the updated classification scheme of Pêgas et al. 2023 (*59*), the Jehol tapejarids are assigned to three taxa: *S. dongi*, *Huaxiadraco corollatus*, *E. lii* (*59*), and the newly published *E. yixianensis* (*60*). The new specimen CUGB-P2201 shares two diagnostic features with *S. dongi*: i, the metatarsal I is almost equal to metatarsal II in length, and longer than metatarsal III, as an autapomorphy of *S. dongi*; ii, the ratio of the length of the fourth wing phalanx to the first wing phalanx in CUGB-P2201 is about 0.27, which is comparable to *S. dongi* but distinct from *E. lii* and *Huaxiadraco corollatus*. The new specimen is also excluded from *E. yixianensis* in having a higher length ratio of the fourth wing phalanx to humerus. In summary, we designate CUGB-P2201 as an individual of *S. dongi*.

**Ontogenetic Status**

The ontogenetic status of pterosaurs is judged primarily by the fusion state of bones and the degree of ossification of the epiphysis (*80*, *81*). There are multiple immature features present in this specimen, i.e. unfused bones: the neural arch and centrum of dorsal vertebrates, scapula and coracoid, distal humeral epiphyses, carpals, extensor tendon process of the first wing phalanx, tibia and proximal tarsals, pelvis. These osteological features suggest this specimen represents a juvenile individual. Most specimens of the tapejarids collected from the Jiufotang Formation are immature individuals, only *S. dongi* D2525 has reached maturity (*65*). The ontogenetic status of this specimen is comparable to the holotype of *S. dongi* and *H. jii* (*56*, *63*), although its humerus has reached 83.4 mm, and the wingspan is over 1500 mm.

**Anatomical terminology**

The anatomical terminology used in the description part of the supplementary information is based on Romerian nomenclature and orientation, using anterior/posterior instead of cranial/caudal (*82*). The orientation regarding the wing elements is based on the previously proposed flight posture (*40*, *83*, *84*).

**Supplementary description of soft tissues**

The specimen CUGB-P2201 preserved three different types of pycnofibres: monofilaments, "brush-like" pycnofibres and "feather-like" pycnofibres. Monofilaments are the simplest filamentous structures without branches (Fig. 1G), they are distributed around the pectoral girdle and scattered dorsal vertebrae. Monofilaments are usually slightly curved and densely packed together. Overlapping of different monofilaments is often observed. Individual monofilament ranges from 0.05-0.20 mm in diameter, and the maximum length preserved is 6.6 mm (actual length should be much longer). Monofilaments have been widely reported in pterosaurs (*11–13*, *16*, *20*, *85–87*), similar unbranched monofilaments are also reported in dinosaurs (*88*, *89*).

The "brush-like" pycnofibres are rare and found only near the pectoral girdle, along with monofilaments. The "brush-like" pycnofibres show a long, thick proximal shaft and a series of branches at the distal end. A total of four pycnofibre of this type were found in the same region (Fig. 1E,F), two of them preserved the distal branches (1, 2 in Fig. 1F) whereas the other two only preserved the proximal shaft (3, 4 in Fig. 1F). The largest one preserved its complete structure (1 in Fig. 1F), its proximal shaft is about 8.5 mm long and 0.72-0.78 mm wide, and the distal branches are about 1.7-2.6 mm in length and 0.17-0.19 mm in diameter, which are of roughly the same diameter as the nearby monofilaments. Another individual preserving the distal branches (2 in Fig. 1F) has a proximal shaft length of 3.0 mm and a width of 0.37-0.47 mm, with the distal branches 0.8-1.3 mm long and about 0.13-0.20 mm wide.

The "brush-like" pycnofibres can be clearly classified as a single type rather than a stack of monofilaments because: i) the "brush-like" pycnofibre 1 in Fig. 1E preserved a complete, long proximal shaft without any trace of monofilaments around it; ii) although the distal branches of "brush-like" pycnofibre 1 is close to the nearby monofilaments, its distal site is clear and no branch exceeds its outline, while the distribution of monofilaments is usually irregular; iii) the "brush-like" pycnofibre 2 also preserves the distal branches while there are no monofilaments around it. "Brush-like" pycnofibres were previously reported in anurognathids CAGS-Z070 (termed type 2 filament) (*12*), in contrast, this type of pycnofibre in CUGB-P2201 possesses a thicker proximal part than in anurognathids (table S2), which allows them to be readily distinguished from the monofilaments.

Three "feather-like" pycnofibres are distributed on the surface of right carpals, these pycnofibres are long and curved, tapering distally, with serial branches on both sides (Fig. 1C). The diameter of the central shaft in these "feather-like" pycnofibres are roughly 0.23-0.41 mm, and the preserved length of the shaft is 4.0-5.1 mm. The branches on either side are about 0.3-0.7 mm long and 0.06-0.08 mm wide. The "feather-like" pycnofibres were previously reported on the occipital crest of *Tupandactylus* cf. *imperator* (*13*) and were proposed to be similar to the stage IIIa of the developmental model for extant feather (*13*, *23*). The "feather-like" pycnofibres here are similar in morphology to those of *Tupandactylus* cf. *imperator* but have longer branches (table S2).

The actinofibrils (structural fibres), a structure supporting and regulating the pterosaur wing membrane, are distributed between the right wing metacarpal and wing digit (fig. S1B-D). Unlike the pycnofibres, these actinofibrils are rigid and extend a long distance. The actinofibrils are usually parallel or subparallel packed with a density of 3-5 fibres / mm. The diameter of a single actinofibril is 0.07-0.22 mm. The complete length of an individual fibre cannot be assessed due to incomplete preservation.

This specimen also preserves the soft tissues associated with the cranial crest. A series of differently oriented crest fibres are preserved on the dorsal premaxillary crest (fig. S1E,I-K), implying the general contour of the head crest. On the ventral side of the premaxillary crest, there are extensive dark soft tissues (fig. S1E-H). Of note is a piece of flaky soft tissue immediately adjacent to the middle position of the premaxillary crest (Fig. 1H, fig. S4), which exceeds the thickness of the surrounding tissues and is distributed in a black and white mottled pattern. The soft tissue on the ventral side of premaxillary crest should represent skin remnant because: 1) there are not any trace of pycnofibres in this region; (2) the single morphology of melanosomes within the soft tissue in this region (fig. S4H) contradicts the different morphologies of melanosomes in monofilaments (fig. S6, described in detail below), thus the possibility of pycnofibre stack can be rule out; (3) these soft tissues are continuously distributed along the ventral side of premaxillary crest, therefore most likely represent remnants of skin.

**Identification of melanosomes within monofilaments and skin remnant**

There is a definite difference in elemental composition between the carbon-rich monofilaments and siliceous sediments (fig. S5). These monofilaments consist of a series of organic debris under SEM, FIB results showed that these debris contain many microbodies and a matrix that encapsulates them with distinct electron density (Fig. 2C,F,I). The size and morphology of these microbodies (ovoid to rod-shaped) is highly similar to melanosomes in extant feathers and hairs. We identified these microbodies as melanosomes preserved in monofilaments because: i, EDS showed these microbodies are mainly composed of C and O, in addition to Ca, Si, Ti and other trace elements (fig. S9A-G), which is definitely different from the minerals in sediments (fig. S9M-O); ii, ToF-SIMS analysis of monofilament revealed organic negative ion signals previously presumed to represent eumelanin (*18*, *90–91*) (green arrowheads in fig. S9U, $C_3N^-$, m/z=50; $C_5N^-$, m/z=74; $C_5NO^-$, m/z=90), which suggests these microbodies in monofilaments are melanosomes containing eumelanin (fig. S9P-U); iii, there is no trace of binary fission and exopolymeric substances (EPS) between microbodies as in bacteria; iv, three-dimensionally preserved microbodies and associated imprints are restricted to monofilaments; v, definitely directional arrangement and organization of microbodies were observed in monofilaments (Fig. 2), whereas decay leads to dispersion of layers or organization (*92*); vi, the internal electron density of these microbodies is uniform, in contrast to the less electron-dense core in bacteria (*9*, *93*), ruling out the possibility of bacteria. On this basis, the matrix enclosing the melanosomes is reminiscent of the keratin matrix in feathers. EDS analyses showed that the matrix contains the elements C, O but high Si content compared to melanosomes (fig. S9H,I), which are distinctly different from minerals (fig. S9N,O). The ToF-SIMS analysis of monofilament showed negative ion signals related to silicate and phosphate (yellow and blue arrowheads in fig. S9U, $SiO_2^-$, m/z=60; $SiO_3^-$, m/z=76; $SiO_3H^-$, m/z=77; $SiAlO_4^-$, m/z=119; $PO_2^-$, m/z=63; $PO_3^-$, m/z=79) except eumelanin (green arrowheads in fig. S9U), which suggests that monofilaments underwent partial silicification and phosphatization after burial, both of which are important modes of soft-tissue fine-structure preservation in fossils (*94*, *95*). Melanosomes are usually wrapped in a keratin matrix in extant feathers or hairs, here we did not find definite ion peaks associated with proteins in positive ion spectra of monofilaments. The above evidence suggests the molecular composition of the matrix of monofilaments has changed during fossilization. The participation of silicification / phosphatization during burial may increase structural stability and ultimately lead to the preservation of the three-dimensional structure. We assumed the matrix retains a volume similar to the original at least within the ordered structure regions (Fig. 2) because: 1. previous fossil reports have demonstrated silicification/phosphatization possesses the potential to preserved high-fidelity, original-scale structures (*94*, *95*); 2. matrix degradation would cause structural collapse, melanosome dispersion and loss of ordered structure; 3. if these regions contain more original matrix, it is improbable that the ordered structure would retain during degradation unless the matrix degradation rate is uniform in all directions, ultimately resulting in a proportionally reduced ordered structure – a scenario difficult to achieve during degradation.

The FIB results revealed the skin remnant on the ventral side of the premaxillary crest also contains many microbodies (Fig. 1H, fig. S4H). These microbodies should also represent melanosomes because: 1) they exhibit similar morphology and size to extant melanosomes; 2) these microbodies possess a uniform internal electron density in contrast to the less electron-dense core in bacteria (*9*, *93*); 3) The ToF-SIMS negative ion spectra of this region also revealed ion peaks associated with eumelanin (green arrowheads in fig. S9V). From the ToF-SIMS results, it is likely the skin remnant near the premaxillary crest underwent a preservation process similar to the monofilaments: the negative ion spectra of the remnant skin also revealed ions associated with silicate and phosphate (yellow and blue arrowheads in fig. S9V). The regional differences in the concentration of melanosomes and the corresponding enrichment of the elements S, Cu, Ti and Ni strongly suggest the melanin-based stripe pattern in the skin here (fig. S4).

**Internal structure of monofilaments**

In addition to the photonic structure and the disordered melanosomes layer underlying it mentioned in the main text, we found more information about the packing structure of melanosomes in the monofilaments through a series of sections. We use the term “Lamina” to refer to the different melanosome organization inside the monofilament. The photonic structure and the disordered melanosomes layer immediately underlying it represent “Lamina-1” and “Lamina-2” respectively.

One of the most complete sections, M2csls1, reveals a continuous three-layer spatial pattern within a single monofilament. In this pattern, the top and bottom layers contain the rod-shaped melanosomes (fig. S6H-L). The melanosomes in the top layer measure 178.02±21.82 nm in diameter (n=270) and tightly arranged, whereas those in the bottom layer are 225.34±28.95 nm in diameter (n=230) and are loosely arranged (L2 and L4 in fig. S6H-L). Sandwiched between both layers are large ovoid and irregularly arranged melanosomes and measure 415.12±77.64 nm in diameter (n=92, L3 in fig. S6H-J). To demonstrate these layering is statistically valid, we performed multimodality test on the melanosome diameter data collected from cross sections at M2csls1 (Data S1, n=475, without any pre-grouping) (*91*). The result indicates melanosome diameter follows a bimodal distribution (log-transformed data: unimodality rejected, $p=0.017<0.05$; bimodality not rejected, $p=0.468$). Subsequently, we fitted Gaussian mixture model to the density distribution of the raw data histogram (*92*), successfully identifying two components separated by a boundary of approximately 300 nm (fig. S12B). We plotted data points with diameters above 300 nm as Component 1 on the cross section, they were found to reside almost entirely within the core region (fig. S12G), confirming this type of melanosomes is primarily distributed in the core region of monofilaments (L3). However, we concluded the other component encompass two potential groups because: (1) after removing data from the core region, the remaining histogram density curve exhibited two potential peaks (red arrowheads in fig. S12C); (2) stratification of melanosomes with varying sizes within 300 nm were observed in collected sections, such as the overlapping region M16ls1 (fig. S8E-G). To verify this, we re-ran multimodality testing and Gaussian mixture model fitting on data from M2csls1 (with Component 1 removed) and M16ls1. Multimodality tests on both data still yield unimodal results (M2csls1: unimodality not rejected, p=0.503; M16ls1: unimodality not rejected, p=0.949), but Gaussian mixture modeling for M16ls1 successfully identified two components (fig. S12E, Component 2 and Component 3). These components exhibit overlapping distributions, explaining their difficulty in detection by multimodality test. We distinguished these two components using 195 nm as the boundary and projected points onto cross sections of M2csls1 and M16ls1, the result revealed distinct regional enrichment of melanosomes for both components, primarily concentrated on different sides of monofilaments (fig. S12F-H). Notably, in M2csls1, Component 2 predominantly localized to the dorsal side of the core region, while Component 3 mainly accumulated on the ventral side of the core region (fig. S12G), confirming the continuous three-layer spatial pattern observed in sections (Component1, 2, 3 correspond to L3, L2, L4). Since the diameter of melanosomes in three layers are quite different, we can link them to the previously described "Lamina-1" and "Lamina-2" by the diameter comparison. The average diameter of the disordered melanosomes immediately under the photonic structure is 175.06±19.62 nm (n=35, "Lamina-2"), this dimension is essentially the same as the top layer in the M2csls1 section (L2 in fig. S6H-L), so we determine that the two are actually the same part of structure, and the three layers of melanosomes in section M2csls1 (L2-L4) correspond to "Lamina-2", "Lamina-3" and "Lamina-4". Taken together, we build an internal model of monofilament based on above results, which should include at least four laminae from dorsal to ventral: (i) photonic structure composed of ordered melanosomes (violet arrow heads in Fig. 2, Table 1); (ii) dorsal rod-shaped melanosomes lamina (the part below the yellow dashed lines in Fig. 2C,F, L2 in fig. S6H-L, Table 1); (iii) a third lamina contains large ovoid melanosomes (Lamina-3 in fig. S6H-J, Table 1); (iv) ventral rod-shaped melanosomes lamina (Lamina-4 in fig. S6H-L, Table 1). The dorsoventral orientation of the reconstructed model is based on: (1), the outer part of the photonic structure (Lamina-1) at position M1ls1 is a smooth surface (Fig. 2B), it should located on the dorsal side to reflect light and form iridescent color; (2), Lamina-3 is located in the central region at section M2csls1, and the large size and morphology of the melanosomes in it are similar to those distributed near the core of non-iridescent structural barbs/gray feathers (*30*), thus we hypothesize that Lamina-3 is located at the center of monofilaments, whereas Lamina1-Lamina2 and Lamina4 are located on the dorsal and ventral sides of the monofilament, respectively.

Similar internal organization was also found in other sections such as M17cs1 and M15ls1 (fig. S6A-G, M-P). The organization of melanosomes in different sections can correspond to our model by the diameter of melanosomes in it. The sections at position M17cs1 contains a series of consecutive cross sections (fig. S6A-F) and a longitudinal section (fig. S6G) of an individual monofilament M17, which showed elongated rod-shaped melanosomes (Lamina-2) overlying larger ovoid melanosomes (Lamina-3), representing the dorsal part of the monofilament. The sections at position M15ls1 show large ovoid melanosomes (Lamina-3) overlying slightly smaller rod-shaped melanosomes (Lamina-4), representing the ventral part of the monofilament (fig. S6M-P). The maximum preserved thickness of Lamina-2, 3 and 4 in different sections is 12.04 μm, 5.96 μm and 17 μm respectively. The width of the monofilaments in the sample region ranged from 75-

183 μm, and the thickness of a complete monofilament is at least 37 μm based on the maximum preserved thickness of different laminae, which should have an elliptical cross section with a height-to-width ratio of 1.97-4.82. The melanosome organization is even preserved in imprints, showing large ovoid melanosome impressions contact with the rod-shaped melanosomes impressions with a diameter of 252.58±28.03 nm (n=23, fig. S7D-F).

To assess the effect of potential monofilament overlays on melanosome organization, we also performed FIB cuts on three regions with obvious two monofilament stacks for comparison. The result showed that the boundaries between two monofilaments consist of only rod-shaped melanosomes (Lamina-2 or Lamina-4), while the central large melanosomes are absent (fig. S8A-I, violet dashed lines). In some overlaid regions, the monofilament below the overlay boundary still preserved the original internal structure (fig. S8A-D, yellow dashed lines). All these results confirm that the inferred laminar order is correct. The absence of ordered photonic structure in the stacked region may be due to perturbation during burial or it was lost before stacking (the difference in diameter between the melanosomes in Lamina-1 and Lamina-2 is smaller compared to other laminae, therefore it would be difficult to distinguish the two once the ordering of Lamina-1 is perturbed and mixed with Lamina-2).

The overlay of ordered melanosomes (Lamina-1) in position M2csls4 by some larger-sized melanosomes (Fig. 2F) represents taphonomic artifact because: (1) other sections in this position show these larger-sized melanosomes are in contact with other melanosomes at an angle (violet dashed lines in fig. S8J-P), these irregular boundaries are different from the near-parallel boundaries representing the original internal structure (e.g. yellow dashed lines in fig. S6H-L, S8A-D), and in particular fig. S8N shows the large-sized melanosomes are extruded and lifted up from the bottom; (2) the diameter of overlying larger-sized melanosomes is 237.41±34.58 nm (n=64), which is almost the same as the melanosomes in Lamina-4 located at the bottom. Therefore, the overlying melanosomes at position M2csls4 represent Lamina-4 originally located at the bottom of monofilaments. Given that there is no obvious monofilament overlay in this position, it is most likely caused by stagger of single monofilament.

The melanosomes in different laminae show difference in morphology and size (fig. S11A, C-E). When compared with the data on melanosomes from extant birds, we found the average morphology of melanosomes in Lamina-1 (photonic structure) and Lamina-2 falls at the edge of the morphospace of cylindrical melanosomes in extant iridescent feathers (fig. S11B), which have shorter lengths and smaller sizes. Noteworthy are the centrally located large melanosomes (Lamina-3), whose morphology falls within the morphospace of melanosomes in extant brown, grey, non-iridescent feathers and penguin feathers (fig. S11B). Melanosomes in brown feathers typically show smaller size, while large melanosomes in penguin could be adaptations to selective pressures on color and material properties (*96*). Melanosomes in non-iridescent structural color feathers are concentrated in the barb core, and those in grey feathers similarly show a greater concentration in the core than the cortex (*30*), this distribution is similar to Lamina-3 located in the core of monofilaments. The tube-like morphology of monofilaments was previously reported in anurognathid pterosaur (*12*), the light-toned axial regions also suggest a potential central cavity in pterosaur monofilaments similar to the medulla of feathers.

**Potential "surface" of monofilaments**

Some sections also show the outermost "surface" of the monofilament. The monofilament portions located at position M1ls1, M2cs5, M17ls1, M24cs1 show a smooth and roughly flat surface (Fig. 2B, fig. S8Q,U,W). These "surface" layers overlie the melanosomes and matrix with a constant thickness (blue arrows in fig. S8R,S,V,X). Based on the arrangement and size of the internal melanosomes, we can determine that M1ls1 and M17ls1 represent the dorsal side of monofilament (fig. S8Q-S, surface over Lamina-1 and Lamin-2), and M2cs5, M24cs1 represent the ventral side of monofilament (fig. S8U-X, surface over Lamina-4). We measured the potential "surface" thickness of the dorsal and ventral sides of the monofilaments to be 381.74±30.15 nm and 283.27±52.28 nm, respectively, based on two sections (M17ls1 and M24cs1). EDS results showed that the "surface" layer is mainly composed of C, O and other mineral elements such as Ca, Si (fig. S9J-L). The distinct carbon peaks in these spectra are different from those of calcite and minerals near monofilaments (fig. S9M-O), supporting an organic origin rather than a mineral layer. The position and morphology of these surface layers are similar to the cortex in extant feather (fig. S8T). To investigate the composition of these surface layers, we performed FIB-ToF-SIMS analysis at position M24cs1 (fig. S8W, S9W). The ToF-SIMS positive ion spectra showed peaks associated with calcium ($Ca^+$, m/Q=40) and calcium-containing organic complexes ($CaCN^+$, m/Q=66, green arrowheads in fig. S9Y), both ions show enrichment in the top "surface" layer (red arrowheads in fig. S9X, the contact relationship of "surface" layer, melanosomes and matrix can be seen in fig. S8X), consistent with the EDS results. The negative ion spectra showed tentative organic negative ion signals compared to the sediment (green arrowheads in fig. S9Z, $C_4^-$, m/Q=48; $C_4H^-$, m/Q=49; $C_3N^-$, m/Q=50; $C_6^-$, m/Q=72). The depth-signal intensity distribution diagram showed the signal intensity of these tentative organic negative ions $C_4^-$, $C_4H^-$ and $C_6^-$ in the "surface" layer is weaker than melanosomes and matrix below it, while the signal intensity of $C_3N^-$ is more obvious (fig. S9X). The above evidence supports an organic origin of these potential "surface".

**Preservation bias in monofilament internal structure and melanosome imprints**

The complex organization and arrangement of melanosomes in pterosaur monofilaments provides a model for studying the preservation potential of integumentary structures in pterosaurs. Our results show that of sixteen monofilaments examined, five preserved at least two laminae of internal melanosome organization. Most of the monofilaments preserved Lamina-2 or Lamina-4 melanosomes internally, whereas the Lamina-1 (photonic structure) was found in only two monofilaments, and the centrally located Lamina-3 was found in only four monofilaments. The differential preservation may be due to: (1), Lamina-2 and Lamina-4 have a greater thickness than Lamina-1 and Lamina-3 (fig. S6Q) and represent the major components of the dorsal and ventral side of the monofilament; (2) Lamina-1 (photonic structure) is located near the dorsal surface with a small thickness, makes it more susceptible to perturbation or degradation during burial; (3) Lamina-1 and Lamina-2 are less morphologically distinct than other laminae (fig. S11B, Table 1), thus would be difficult to recognize if the order of Lamina-1 was disturbed and mixed with Lamina-2; (4) as mentioned above, Lamina-3 may be located in the center of the monofilaments surrounding a cavity similar to the medulla of feathers, where may be more susceptible to collapse and degradation. It is noteworthy that the melanosome imprints near monofilaments are dominated by Lamina-4 ventrally and Lamina-3 centrally (see below for a discussion of melanosome shrinkage), whereas dorsal Lamina-1 and Lamina-2 are barely preserved in the imprints. This bias preservation in imprints suggests the need for caution when recovering the color of integumentary structure based on melanosome imprints in non-avian dinosaurs, pterosaurs and other archosaurs, where complex melanosome organization may be present.

**Shrinkage of melanosomes during diagenesis**

Previous taphonomic experiments have pointed to the shrinkage of melanosomes under high temperatures and pressures (*18*, *19*), which may affect the color recovered through melanosomes. The melanosome imprints likely formed earlier and reflect the original size more closely (*97*). To exclude the effect of shrinkage, we compared the melanosome imprints with those preserved in three dimensions. The melanosome imprints in the monofilaments region are dominated by rod-shaped imprints and large ovoid imprints (fig. S7A-F, correspond to Lamina-4 and Lamina-3, respectively), Since the melanosomes of lamina-3 are high variable in size (Table 1, table S3) and more susceptible to limited measurements, we selected the rod-shaped imprints of lamina-4 for comparison. Measurements showed that the average size of 3D-preserved melanosomes in FIB sections shrink by 4.49% - 7.63% compared to the imprints (table S3), we therefore estimate the degree of shrinkage of melanosomes in monofilaments does not exceed 10%.

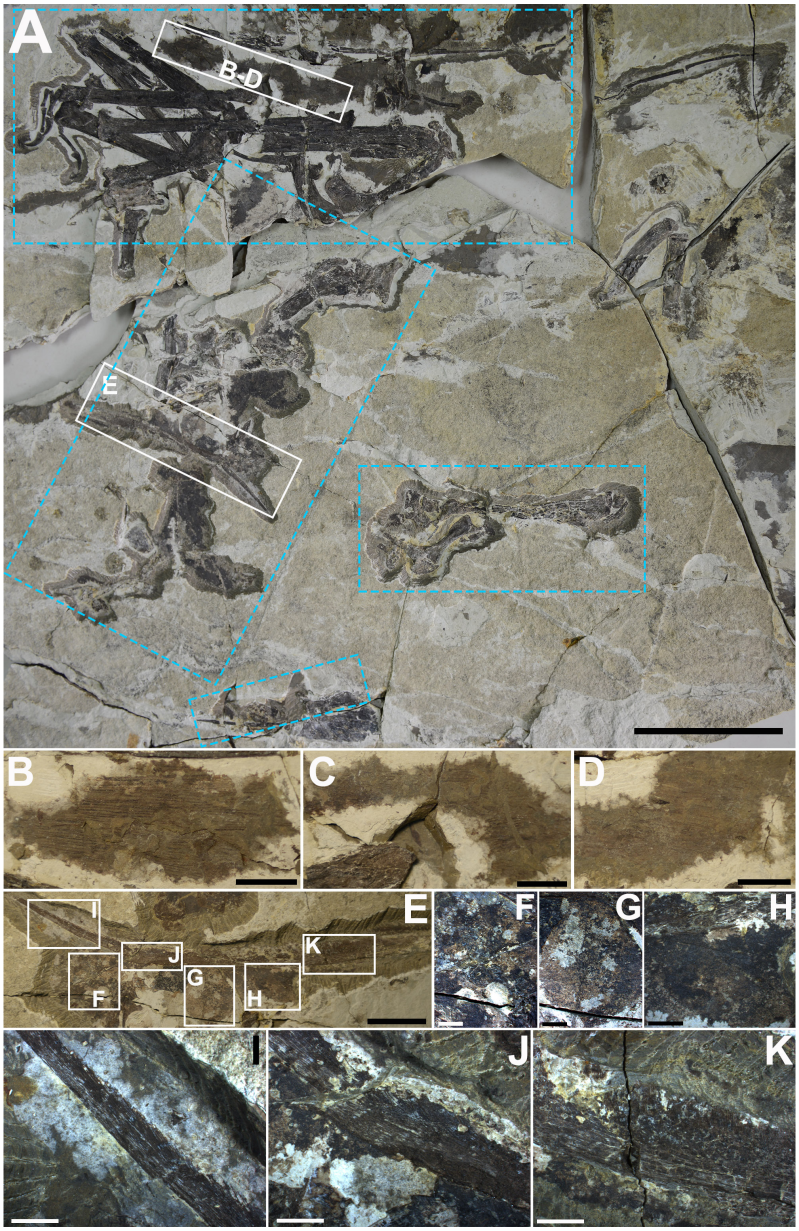


**Fig. S1. Supplementary images of soft tissue in different regions of CUGB-P2201.** (**A**), Overall image of the fossil and regions of soft tissue distribution, four areas for XRF analysis are marked with blue dashed lines. (**B** to **D**), Actinofibrils between the right metacarpal and wing digit. (**E**), Close-up of the premaxillary crest. (**F** to **H**), Soft tissue distribution in (E). (**I** to **K**), Close-up of the crest fibres in different regions, note the change in fibre direction. (**I**), Crest fibres in the posterior region, show a nearly parallel orientation to the bony element. (**J**), Crest fibres in the mid-posterior region, at roughly 30° to the bony element. (**K**), Crest fibres in the anterior region, at roughly 70° to the bony element. Scale bars, 10 cm (A); 2 cm (E); 1 cm (B to D); 4 mm (F to K).

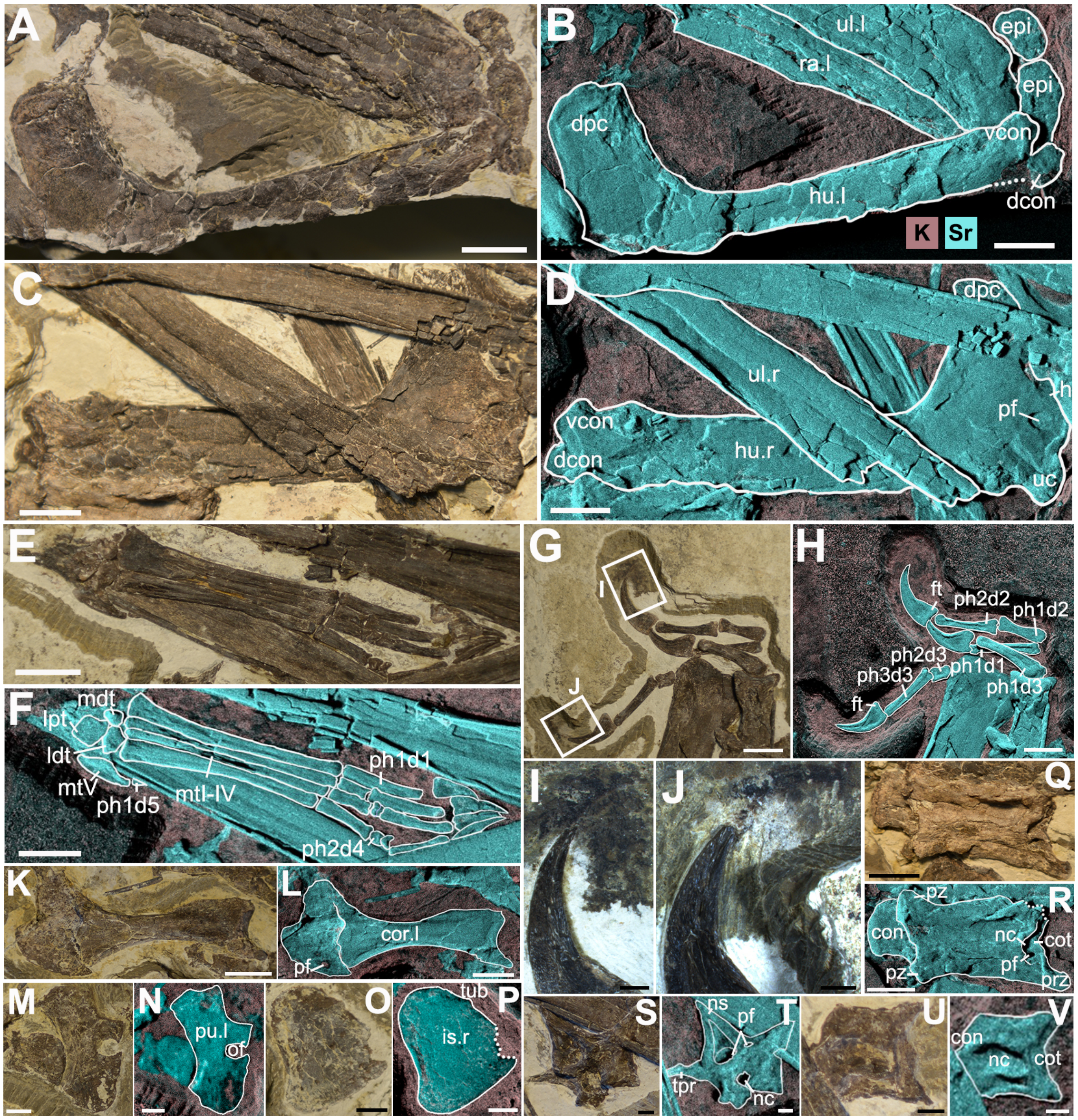


**Fig. S2. Anatomical details of CUGB-P2201.** (**A**), The left humerus. (**C**), The right humerus. (**E**), The right pes. (**G**), The left manus. (**I** to **J**), Close-up of the claw sheath in (G). (**K**), The left coracoid in dorsal view. (**M**, **O**), Part of the pelvic girdle. (**Q**), Isolated cervical vertebra. (**S**, **U**), Isolated neural arch (**S**) and centrum (**U**) of dorsal vertebrae. (**B**, **D**, **F**, **H**, **L**, **N**, **P**, **R**, **T**, **V**), XRF false-color line drawings of corresponding bones, showing the distribution of bone-related element strontium (Sr) and potassium (K) in matrix. Abbreviations: con, condyle; cor, coracoid; cot, cotyle; d, digit; dcon, dorsal condyle; dpc, humeral deltopectoral crest; epi, humeral epiphysis; ft, flexor tubercle; h, humeral head; hu, humerus; is, ischium; ldt, lateral distal tarsal; lpt, lateral proximal tarsal; mdt, medial distal tarsal; mt, metatarsal; nc, neural canal; ns, neural spine; of, obturator foramen; pf, pneumatic foramen; ph, phalanx; prz, prezygapophysis; pu, pubis; pz, postzygapophysis; ra, radius; tpr, transverse process; tub, tubercle; uc, ulnar crest; ul, ulna; vcon, ventral condyle; l, left; r, right. Scale bars, 1 cm (A to H, K to L, Q to R); 5 mm (M to P); 2 mm (I to J, S to V).

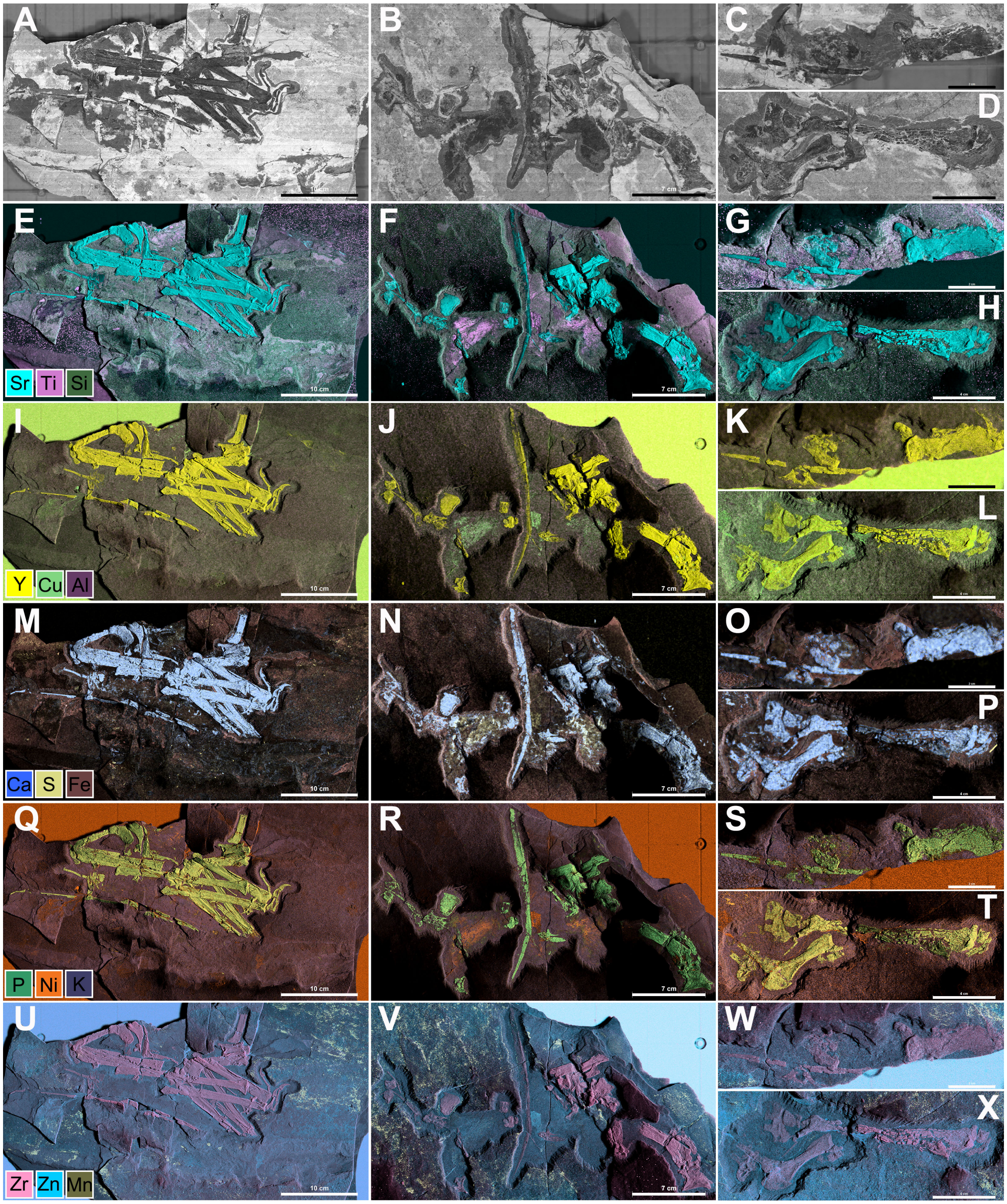


**Fig. S3. Overall element distribution of CUGB-P2201.** (**A** to **D**), Selected scanning regions. (**E** to **H**), Micro-XRF elemental distribution map (false-color images) of Sr-Kα, Ti-Kα, Si-K. (**I** to **L**), Micro-XRF elemental distribution map (false-color images) of Y-Kα, Cu-Kα, Al-K. (**M** to **P**), Micro-XRF elemental distribution map (false-color images) of Ca-Kα, S-Kα, Fe-Kα. (**Q** to **T**), Micro-XRF elemental distribution map (false-color images) of P-Kα, Ni-Kα, K-Kα. (**U** to **X**), Micro-XRF elemental distribution map (false-color images) of Zr-Kα, Zn-Kα, Mn-Kα. Scale bars,

10 cm (A, E, I, M, Q, U); 7 cm(B, F, J, N, R, V); 4 cm (D, H, L, P, T, X); 2 cm (C, G, K, O, S, W).

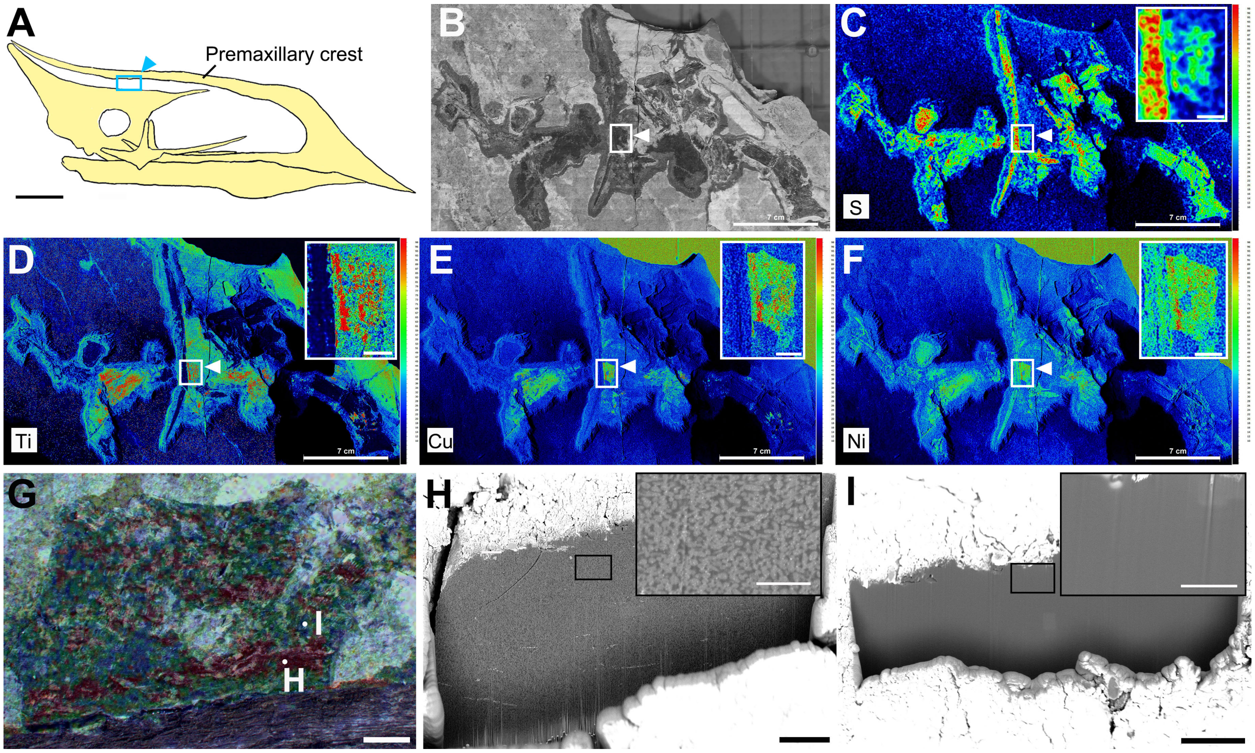


**Fig. S4. Distribution of elements associated with melanosomes in the skin near premaxillary crest.** (**A**), Approximate location of skin remnant on the ventral margin of the premaxillary crest, line drawing is based on the skull of *Sinopterus dongi* holotype IVPP V 13363 (*56*). (**B**), Photograph of the soft tissue-enriched region in CUGB-P2201, the arrow points to the skin remnant. (**C** to **F**), The heat map of elements S, Ti, Cu, Ni, highlighting the distribution and enrichment of individual element. Note the enlarged close-up of flaky skin remnant in (C to F) showing a striated distribution. (**G**), The skin remnant near premaxillary crest overlaid with heat map of Ti, sampling locations are labeled. (**H**), FIB section of the Ti-rich striped region, showing a large number of melanosomes. (**I**), FIB section of the region between Ti-enriched ribbons without melanosomes. Scale bars, 7 cm (B to F); 2 mm (G); 10 µm (H to I); inset, 5 mm (C to F); 2µm (H to I).

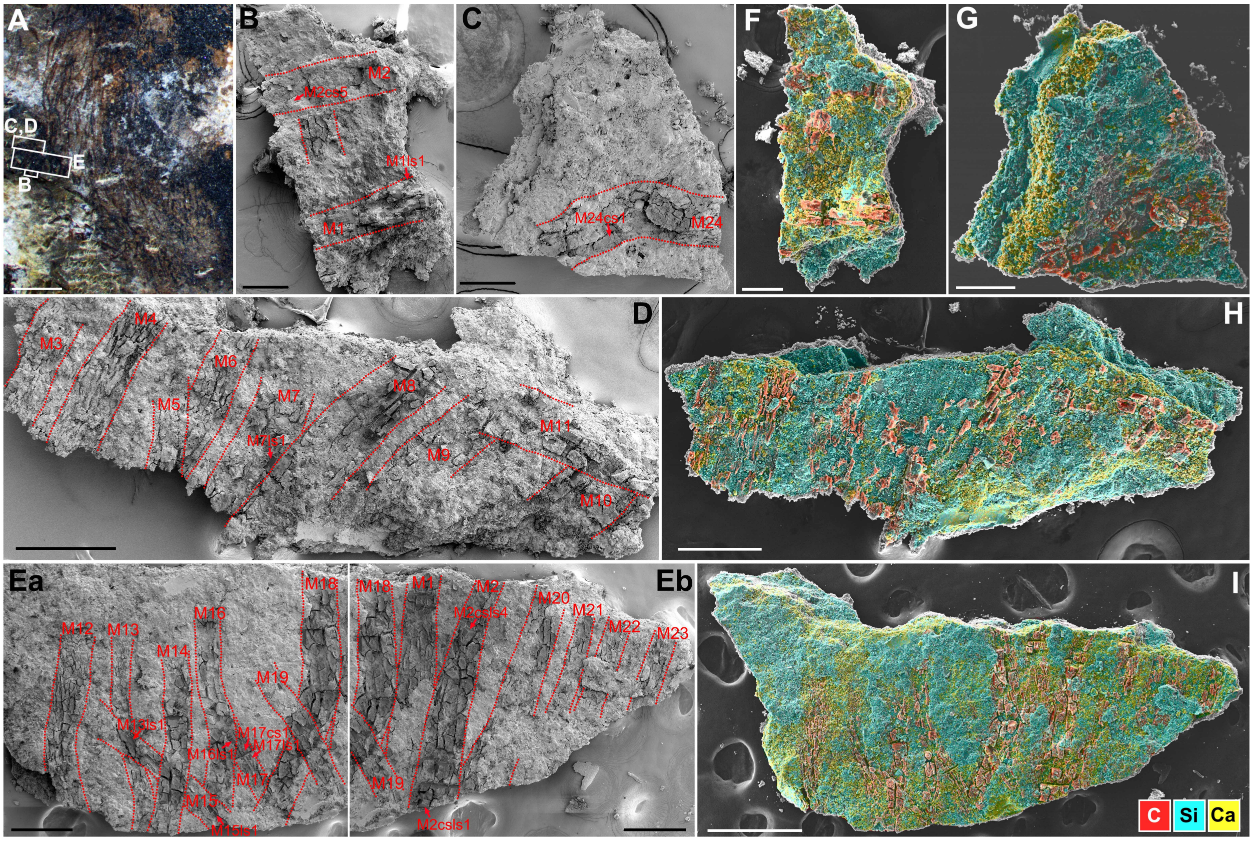


**Fig. S5. Sample location of monofilaments and surface elemental distribution.** (**A**), Sample locations. (**B** to **Eb**), SEM image of sample S1(**B**), S2(**Ea**, **Eb**), S3(**C**), S4(**D**), the contours of monofilaments are marked in figures, FIB sections were made in monofilament M1, M2, M7, M8, M10, M11, M13-M21, M24, the positions of important sections are labeled in figures. (**F** to **I**), EDS false-color images of sample S1-S4 showing the surface distribution of carbon (C, red), silicon (Si, cyan) and calcium (Ca, yellow). Abbreviation: M, monofilament; cs, cross section; ls, longitudinal section. Scale bars, 2 mm (A); 500 µm (I); 250 µm (D, Ea, Eb, H); 100 µm (B, C, F, G).

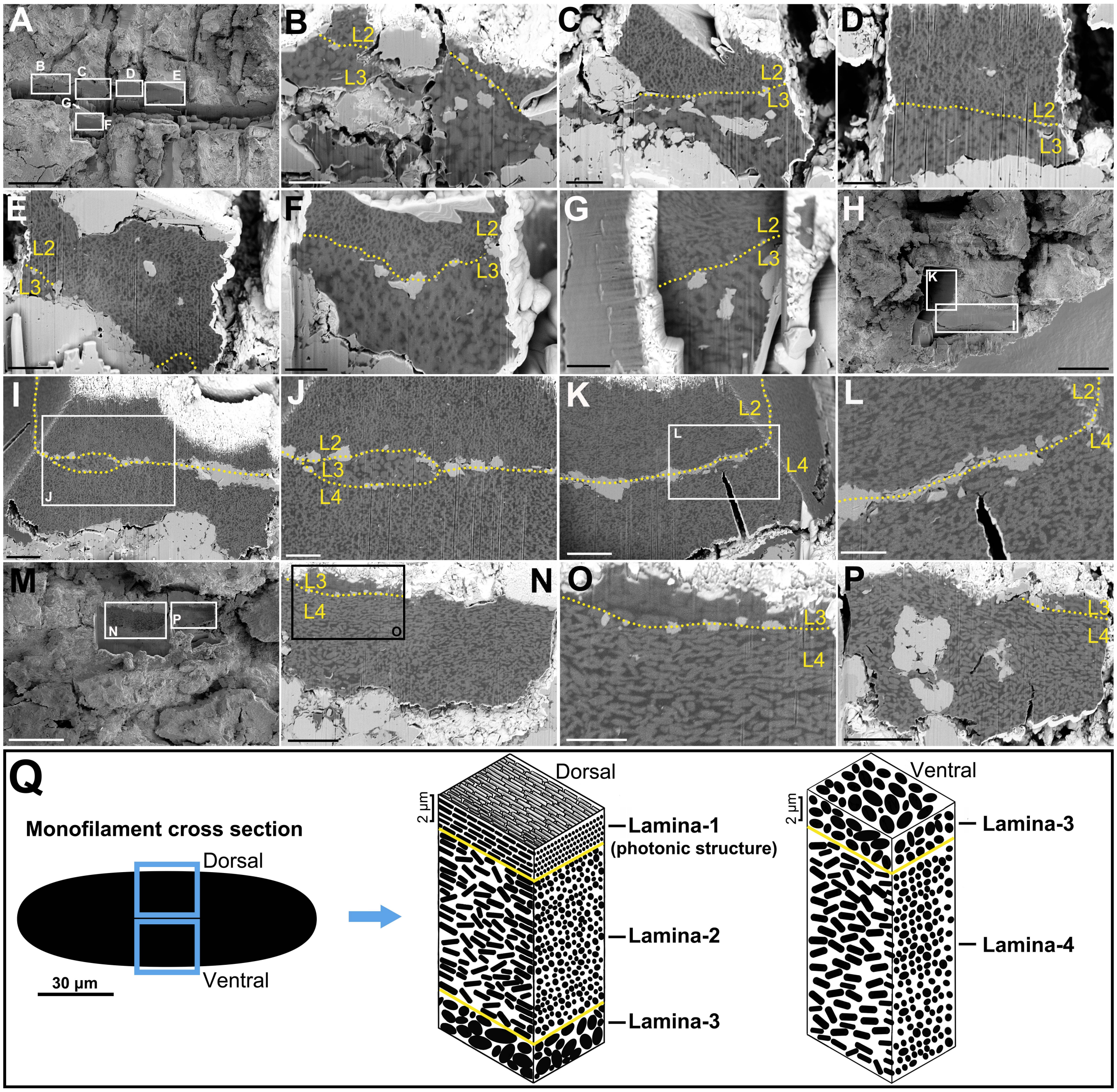


**Fig. S6. FIB sections of the internal structure in monofilaments without overlapping.** (**A** to **G**), Cross sections (**A** to **F**) and longitudinal section (**G**) of M17 at position M17cs1, the sections show a continuous boundary between dorsal rod-shaped melanosomes (L2) and larger central oval melanosomes (L3). (**H** to **L**), Cross section (**H** to **J**) and longitudinal section(**K** to **L**) of M2 at position M2csls1, the cross section shows a complete structure including dorsal and ventral rod-shaped melanosomes with different diameters (L2, L4) as well as large ovoid melanosomes in the central (L3). (**M** to **P**), longitudinal sections of M15 at position M15ls1, showing the boundary between ventral rod-shaped (L4) melanosomes and larger central oval melanosomes (L3). The boundaries of internal structures are marked in figures with yellow dashed lines. (**Q**), Brief reconstruction model of monofilaments in *Sinopterus dongi*, the morphology and size of melanosomes are not drawn to scale. L1-L4: Lamina1-Lamina4. Secondary electron images (A, H, M), backscattered electron images (B to G, I to L, N to P). Scale bars, 25 µm (H, M); 20 µm (A); 5 µm (I, K, N, P); 3 µm (E); 2.5 µm (J, O); 2 µm (B to D, F to G, L).

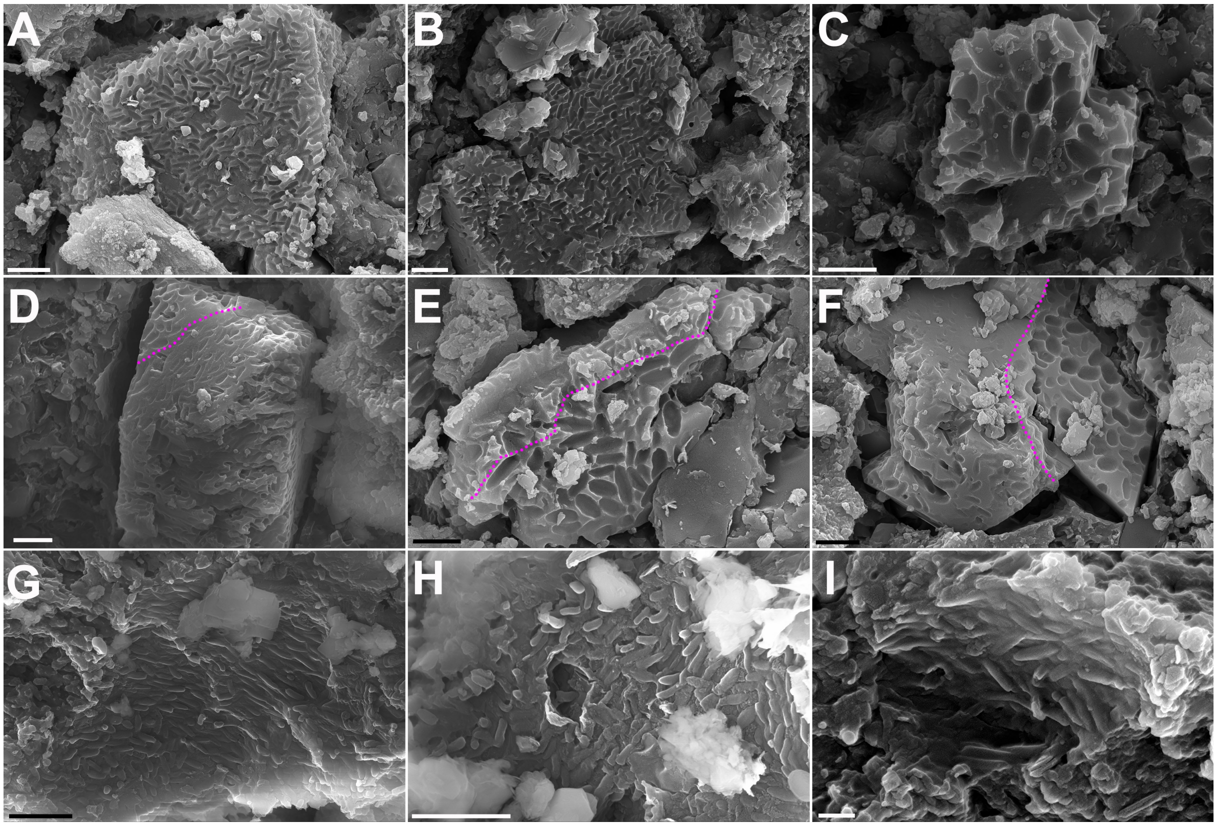


**Fig. S7. Imprints and entities of different shapes of melanosomes observed in monofilaments.** (**A**, **B**), Imprints of rod-shaped melanosomes. (**C**), Imprints of large ovoid-shaped melanosomes. (**D** to **F**), Boundary between the imprints of rod-shaped melanosomes and large ovoid melanosomes, the boundary is marked with violet dashed lines. (**G** to **I**), Three-dimensional preservation of rod-shaped melanosomes with different aspect ratios. Scale bars, 2 μm (A to H); 400 nm (I).

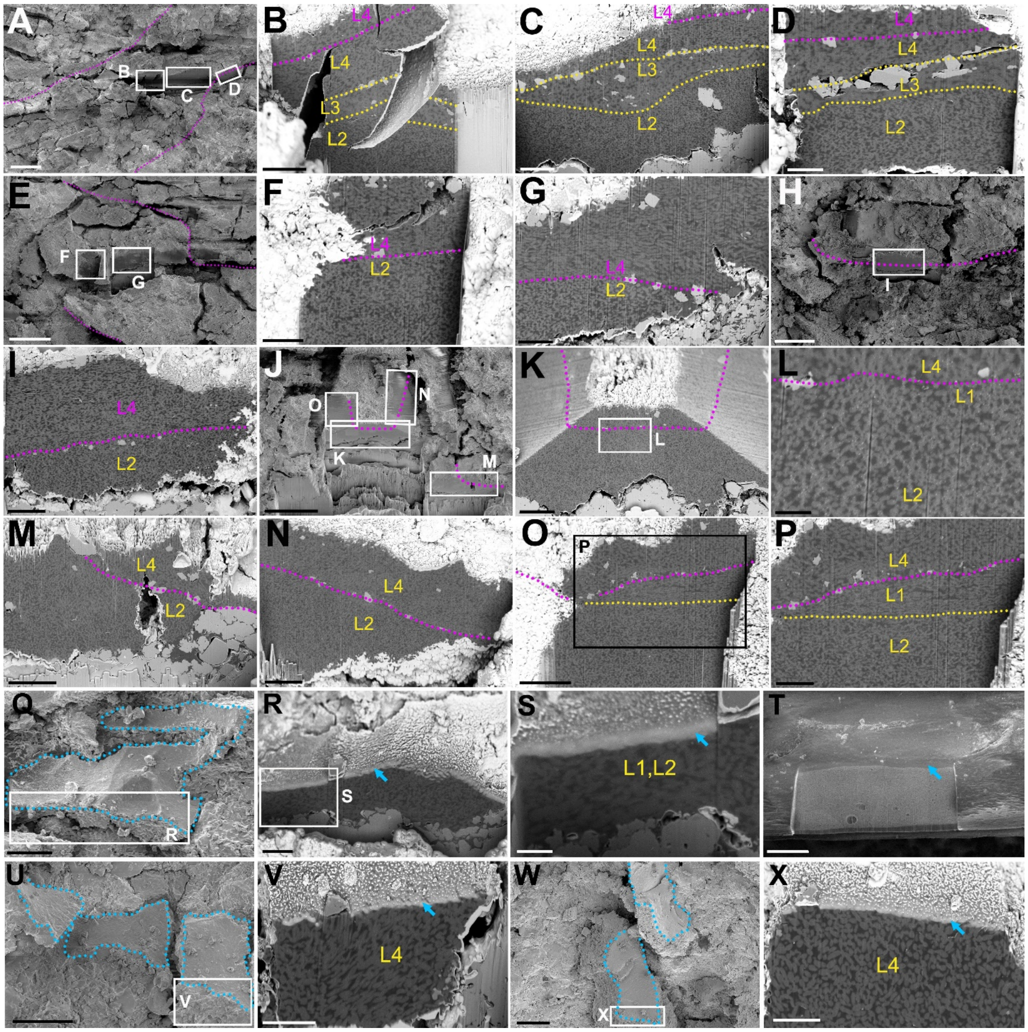


**Fig. S8. FIB sections of different monofilament overlay regions, single monofilament staggered region and "surface" region.** (**A** to **I**), Overlap region and corresponding FIB sections of M13 and M15 at position M13ls1 (**A** to **D**), M16 and M17 at position M16ls1. (**E** to **G**), M7 and M8 at position M7ls1 (**H** to **I**), the stacked M13 showed a continuous three-lamina structure (B to D, yellow dashed lines), the overlay boundaries of different monofilaments are marked with violet dash lines. (**J** to **P**), M2 at position M2csls4 (**J**) and corresponding cross sections (**K** to **M**), longitudinal sections (**N** to **P**). The staggered slip of M2 caused the bottom L4 to overlay the top L1, the overlapping boundaries are marked with violet dashed lines. (**Q** to **X**), Smooth "surface" in fossil monofilament versus extant feather, (**Q** to **S**, **U** to **V**, **W** to **X** are images and sections at position M17ls1, M2cs5, M24cs1 respectively; **T** is the longitudinal section of barb in eastern bluebird *Sialia sialis*, blue dashed lines and arrows mark the surface of monofilament with constant thickness compared to extant barb. L1-L4: Lamina1-Lamina4, laminae of different monofilaments are labelled in different colors. Secondary electron images (A, E, H, J, Q, T, U, W), backscattered electron images (B to D, F to G, I, K to P, R to S, V, X). Scale bars, 25 µm (A, H, J); 20 µm (E); 10 µm (T, U, W); 5 µm (B, C, I, K, M to O, Q); 2.5 µm (D, F, G, P, R, V, X); 1 µm (L, S).

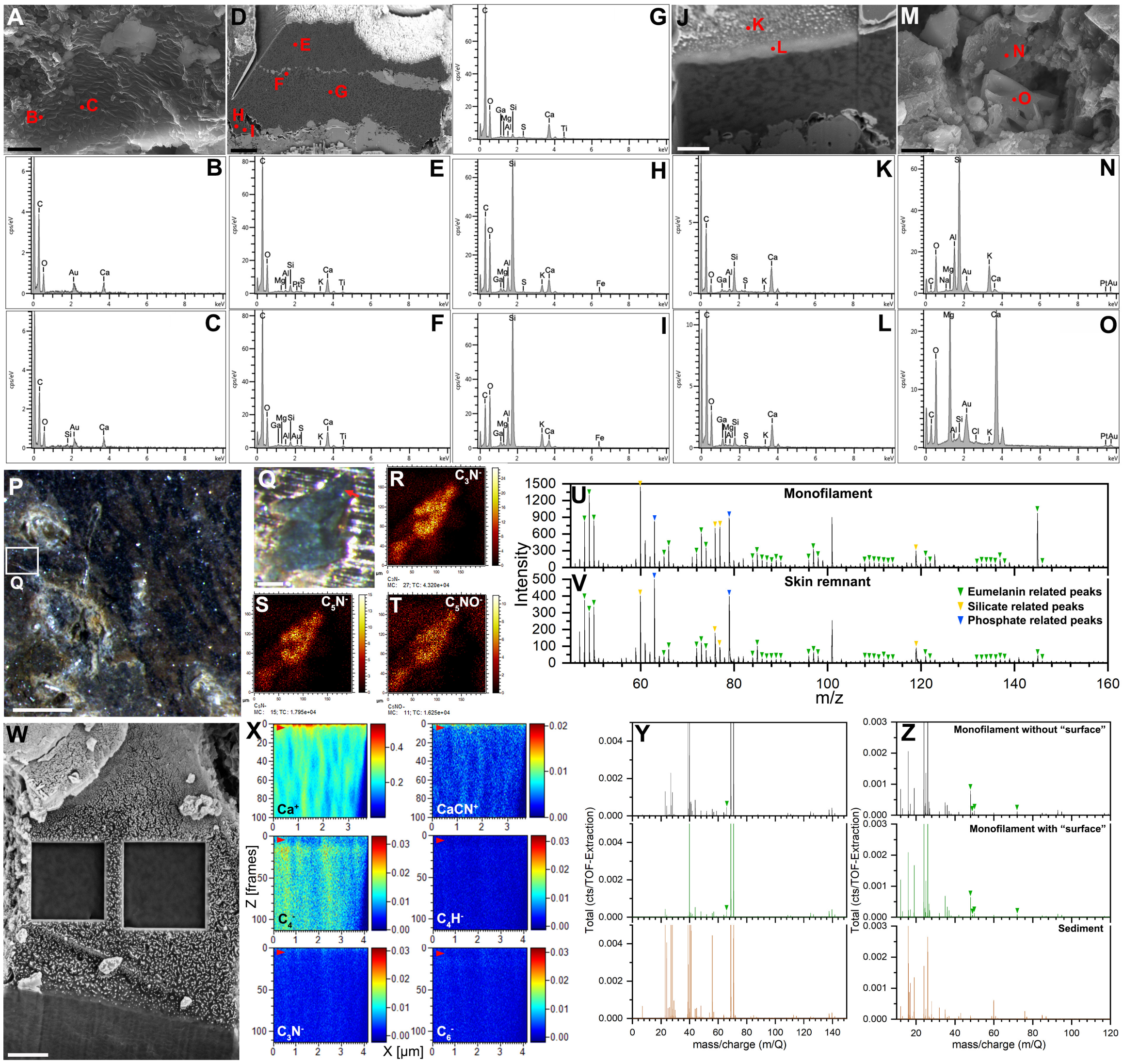


**Fig. S9. EDS and Tof-SIMS of melanosomes, matrix, "surface" and sediments in monofilaments.** (**A** to **O**), EDS of melanosomes in natural sections (**A** to **C**), FIB section (**D** to **I**), potential "surface" at position M17ls1 (**J** to **L**) and minerals (**M** to **O**), (H, I) are regions of matrix without melanosomes. (**P** to **Q**), Sampling position for ToF-SIMS analysis, the red arrow in (Q) points to monofilament. (**R** to **T**), Signal intensity distribution of tentative eumelanin-related ions, showing the correlation with monofilament. (**U** to **V**), ToF-SIMS negative ion spectra of monofilament (**U**) and skin near the premaxillary crest (**V**), the green, yellow and blue arrowheads are tentative eumelanin, silicate and phosphate related ions. (**W** to **Z**), FIB-ToF-SIMS analysis of monofilament "surface". (**W**), Analysis regions at position M24cs1. (**X**), Depth-signal intensity distribution of selected ions, the red arrowheads point to the "surface" at the top. (**Y** to **Z**), Comparison of positive (**Y**) and negative ion (**Z**) spectra of monofilament, "surface" and sediment, the green arrows point to the tentative organic ion peaks. Secondary electron images (A, M, W), backscattered images (D, J). Scale bars, 1 mm (P); 100 μm (Q); 5 μm (D, M); 2 μm (A, W); 1 μm (J).

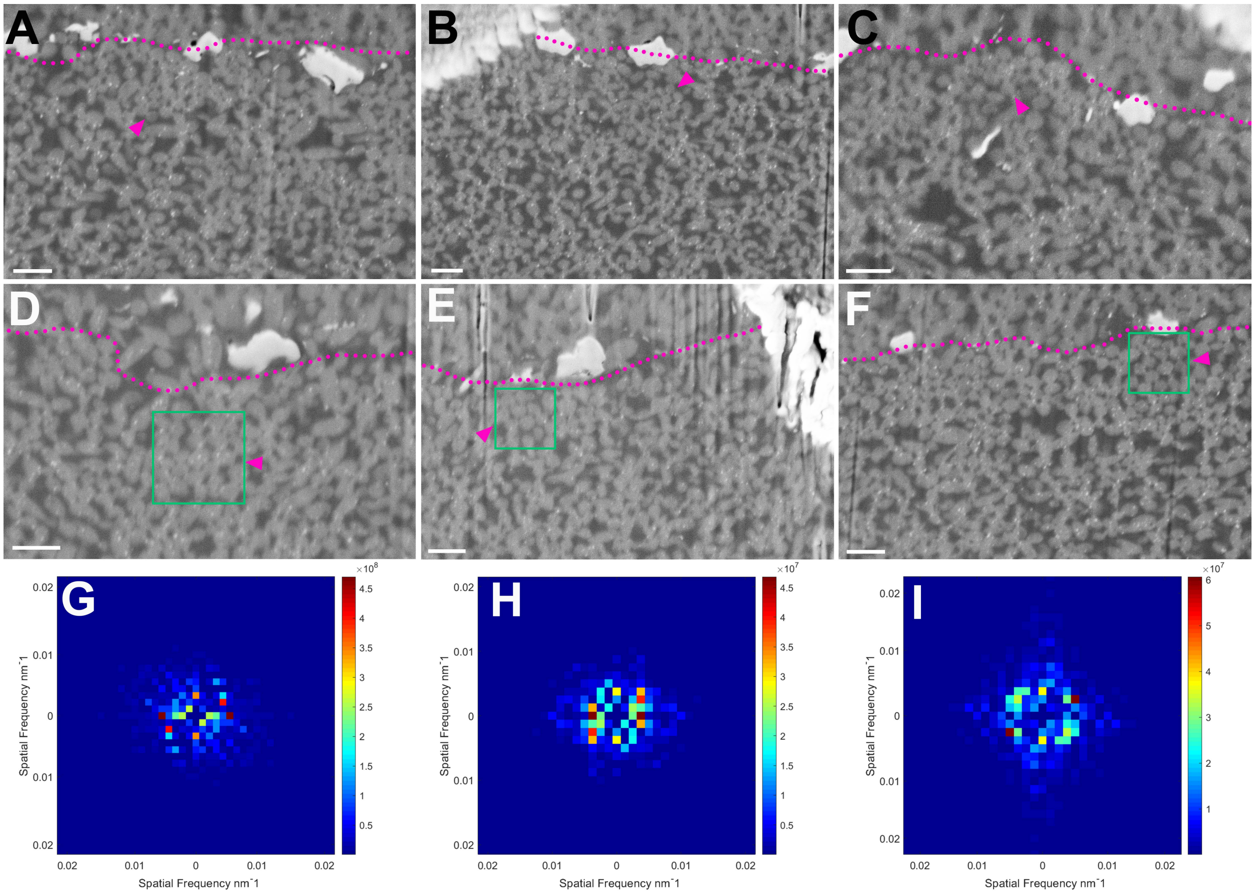


**Fig. S10. Supplementary images detailing the photonic structures in monofilaments.** (**A** to **F**), cross sections of the monofilament at position M2csls4 corresponding to Fig. 2F. The violet arrowheads point to the region where melanosomes are densely packed and the violet dashed line indicates overlapping boundaries caused by staggered slip of monofilament. The green boxes in D-F represent the regions selected for Fourier analysis. (**G** to **I**), 2-D Fourier power spectra of selected regions in D-F, displaying a near-hexagonal pattern. Scale bars, 500 nm (A to F)

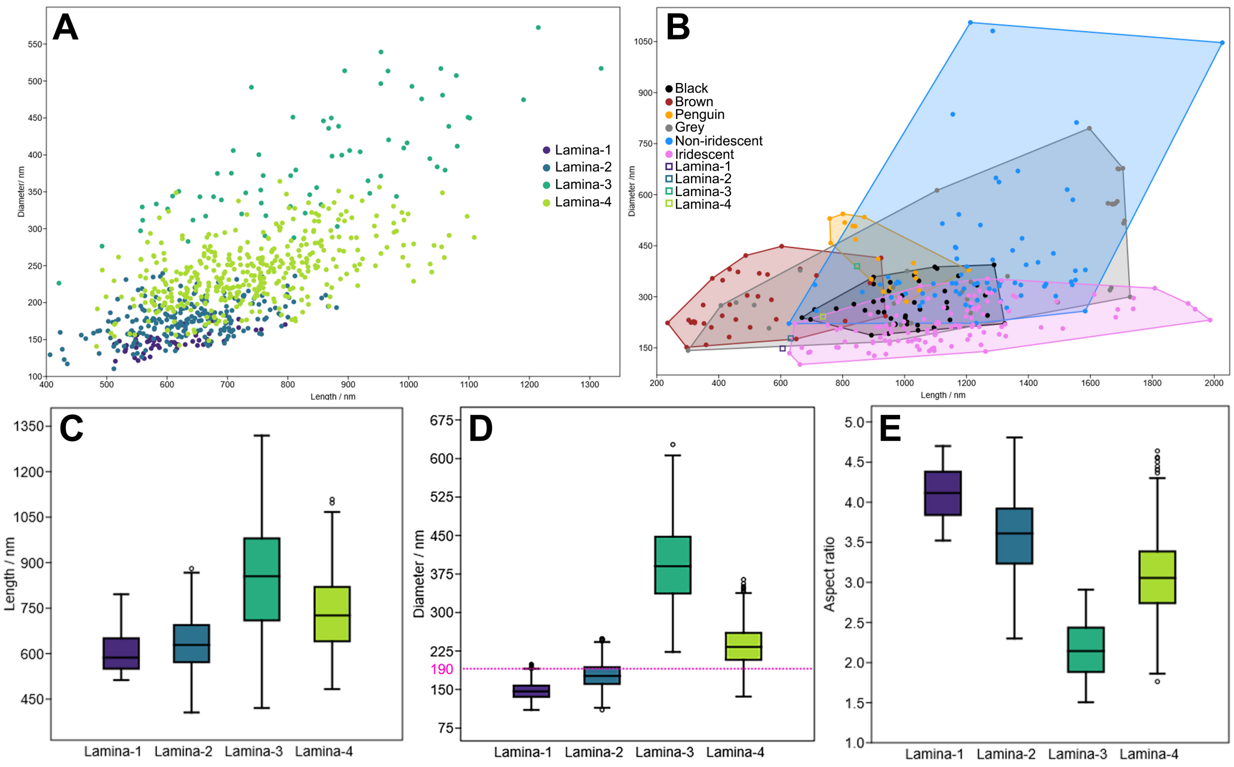


**Fig. S11. Comparison of morphological data between different laminae of melanosomes in monofilaments.** (**A**), Scatter plot of melanosome length (long axis) and diameter (short axis) for different laminae. (**B**), Comparison of melanosomes in different laminae with those in the feathers of extant bird, data on melanosomes in extant feathers were obtained from published literature. (**C** to **E**), Box plots of length (**C**), diameter (**D**) and aspect ratio (**E**) for different laminae of melanosomes. The violet dashed line in D highlight the 190 nm boundary line of the “thin melanin layers” proposed by Nordén et al., 2021.

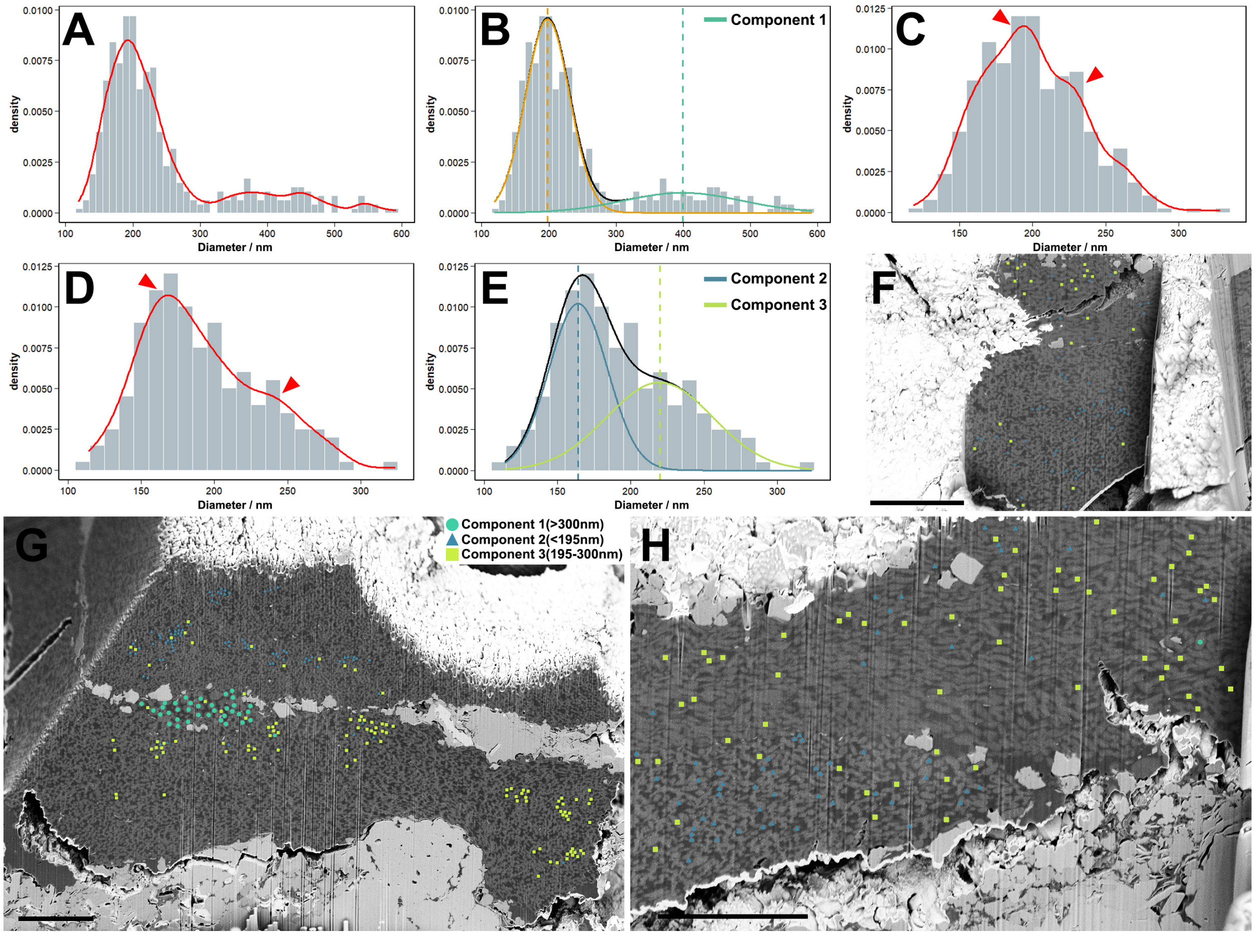


**Fig. S12. Statistical analysis of internal melanosome structure within monofilaments.** (**A**), Density histogram with kernel density curve for melanosome diameter data in M2csls1 cross sections. (**B**), Gaussian mixture model fitting result of (A). (**C**), Density histogram with kernel density curve for melanosome diameter data in M2csls1 cross sections (data from core region removed). (**D**), Density histogram with kernel density curve for melanosome diameter data in M16ls1, red arrowheads point to two potential peaks. (**E**), Gaussian mixture model fitting result of (D). (**F-H**), M16ls1(F, H) and M2csls1(G) sections after data projection of different components. Scale bars, 5 µm (F-H).

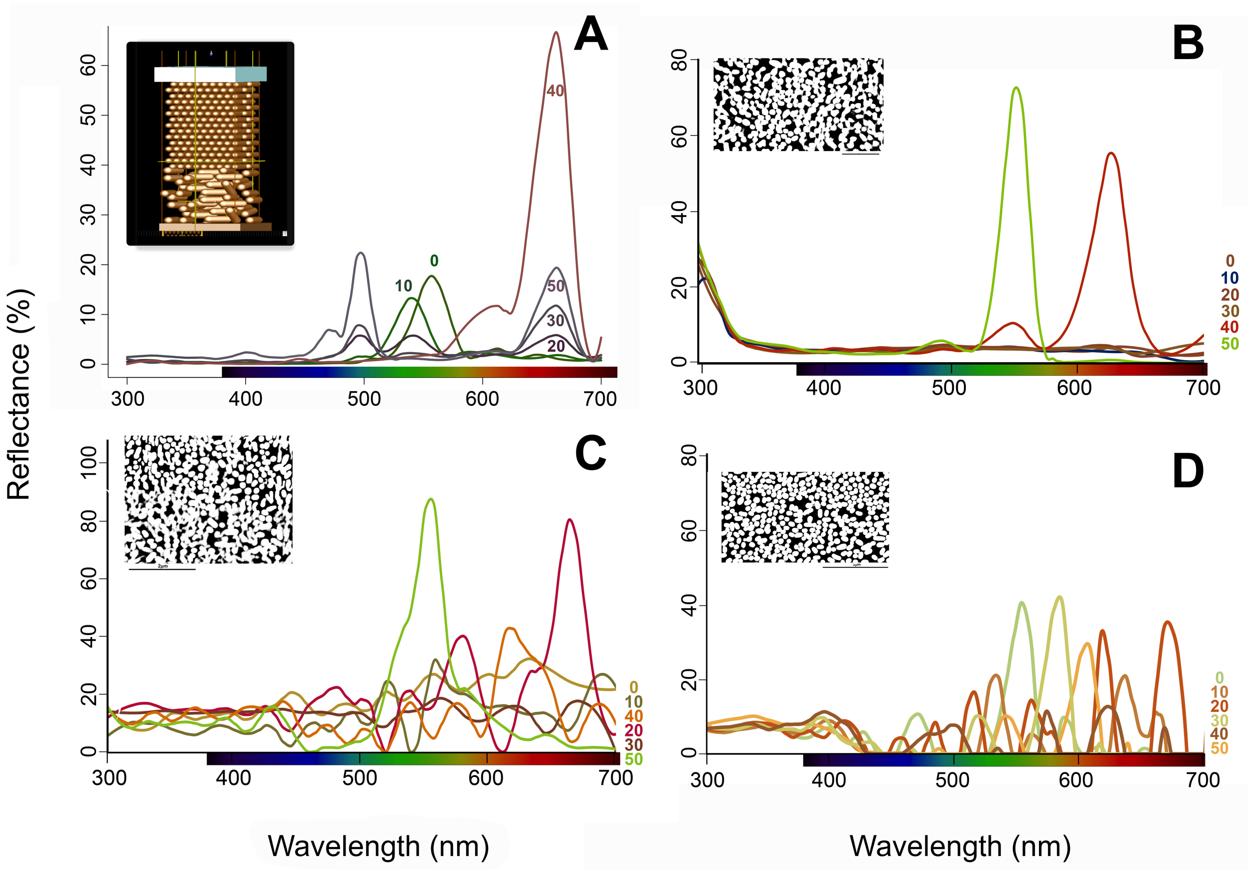


**Fig. S13. Supplementary simulated reflectance from models of idealized and realistic melanosome structures in monofilaments.** (**A**) Idealized model with keratin cortex, expanded melanosomes (accounting for 10% taphonomic shrinkage) and keratin background. (**B-D**) Realistic model based on fossil sections shown in fig. S10D (B), fig. S10E (C), fig. S10F (D), modeled with keratin background but no keratin cortex. Scale bars, 2 µm (C-D); 1 µm (B).

**Table S1. Measurement (in mm) of the postcranial bones of *Sinopterus dongi* (CUGB-P2201).**

| Element | Left | Right |
|---|---|---|
| Scapula | 46.07[p] | 62.20[p] |
| Coracoid | 55.53 | - |
| Humerus | 78.01[p] | 83.42 |
| Ulna | 130.20[e] | 111.70[p] |
| Radius | 125.28[e] | - |
| Pteroid | - | 65.80 |
| Wing metacarpal | 141.06[e] | 103.28[p] |
| Metacarpals I | 28.77[p] | - |
| Manual digit I | -, - | 19.77, 12.93 |
| Manual digit II | -, -, - | 12.66, 16.57, 13.72 |
| Manual digit III | -, -, -, - | 18.81, 4.94, 15.62, 12.04 |

| | | |
|---|---|---|
| First phalanx of wing digit | 174.09 | 97.96[p] |
| Second phalanx of wing digit | 94.35[p] | 114.44[p] |
| Third phalanx of wing digit | 14.11[p] | 76.96[p] |
| Fourth phalanx of wing digit | - | 46.60 |
| Femur | 69.10 | 110.14[b] |
| Tibia | 55.79[p] | 135.48[p] |
| Fibula | 45.72[p] | 15.53[p] |
| Metatarsal I-IV | -, -, -, - | 35.23, 35.11, 33.67, 32.20 |
| Metatarsal V | - | 9.43 |
| Pedal digit I | -, - | 12.99, 7.31 |
| Pedal digit II | -, -, - | 4.76, 12.41, 8.92 |
| Pedal digit III | -, -, -, - | 6.73, 1.67, 11.43, 8.01 |
| Pedal digit IV | -, -, -, -, - | 9.55, 2.03, 1.71, 9.72, 9.70 |
| Pedal digit V | - | 2.53 |

[p], Preserved length; [e], Approximate estimated length (from impressions in surrounding rock); [b], False length due to severe compression and fragmentation; -, not preserved.

### Table S2. Morphological data for differnet types of pycnofibres.

| Pycnofibre morphology | Branching | Taxon | Shaft / mm | Branch / mm | Distribution region | Potential function |
|---|---|---|---|---|---|---|
| 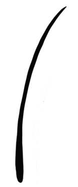 | none | *Sinopterus dongi* (CUGB-P2201) | 6.6* (length) 0.05-0.20 (width) | N/A | pectoral girdle, scattered dorsal vertebrae | thermal insulation, display |
| | | *Tupandactylus* cf. *imperator* (MCT.R.1884) | 30.0 (length) 0.06-0.09 (width) | N/A | proximal portion of the occipital process | |
| | | anurognathids (CAGS–Z070 & NJU–57003) | 3.5-12.8 (length) 0.07-0.43 (width) | N/A | head, neck, shoulder, torso, all four limbs and tail | |
| 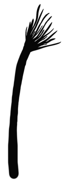 | terminal | *Sinopterus dongi* (CUGB-P2201) | 3.0-8.5 (length) 0.37-0.78 (width) | 0.8-2.6 (length) 0.13-0.20 (width) | pectoral girdle | tactile sensing |
| | | anurognathids (CAGS–Z070) | 2.0-13.8 (length) 0.08-0.18 (width) | 0.05-0.07(width) | neck, proximal forelimb, plantar metatarsus and proximal tail regions | |
| 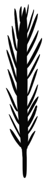 | along the shaft | *Sinopterus dongi* (CUGB-P2201) | 4.0-5.1* (length) 0.23-0.41 (width) | 0.3-0.7(length) 0.06-0.08 (width) | carpal | thermal insulation, display |
| | | *Tupandactylus* cf. *imperator* (MCT.R.1884) | 2.0-5.0 (length) 0.06 (width) | 0.1-0.2 (length) | distal portion of the occipital process | |
| 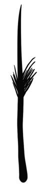 | mid-point | anurognathids (CAGS–Z070) | 4.5-7.0 (length) 0.05-0.45 (width) | 0.02-0.04 (width) | cranium | tactile sensing |
| 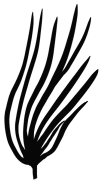 | base | anurognathids (CAGS–Z070 & NJU–57003) | N/A | 2.5-8.0 (length) 0.07-0.13 (width) | wing membrane | thermal insulation |

*, preserved length; Gray cells represent data from this specimen (CUGP-P2201).
Data for anurognathids (CAGS-Z070 & NJU-57003) and *Tupandactylus* cf. *imperator* (MCT.R.1884) are cited from Yang et al., 2019 and Cincotta et al., 2022, respectively.

**Table S3. Comparison of three-dimensionally preserved melanosomes in FIB sections with melanosome imprints.**

| Lamina | n | Long axis / nm | Short axis / nm | Aspect ratio |
|---|---|---|---|---|
| **Imprints** | | | | |
| **3** | 218 | 972.06 ± 306.50 | 425.51 ± 141.96 | 2.33 ± 0.43 |
| **4** | 628 | 771.37 ± 129.32 | 256.00 ± 39.76 | 3.05 ± 0.52 |
| **Longitudinal sections** | | | | |
| **3** | 71 | 846.66 ± 183.19 | 389.58 ± 73.60 | 2.18 ± 0.33 |
| **4** | 398 | 736.77 ± 123.81 (4.49%) | 241.90 ± 43.54 (5.51%) | 3.10 ± 0.52 |
| **Longitudinal sections & cross sections** | | | | |
| **3** | 319 | - | 396.34 ± 80.02 | - |
| **4** | 1089 | - | 236.48 ± 40.04 (7.63%) | - |

Brackets indicate the extent of shrinkage of the three-dimensionally preserved melanosomes compared to the imprints.

**Data S1. (separate file)**
